# ExoS Effector in *Pseudomonas aeruginosa* Hyperactive Type III Secretion System Mutant Promotes Enhanced Plasma Membrane Rupture in Neutrophils

**DOI:** 10.1101/2024.01.24.577040

**Authors:** Arianna D. Reuven, Bethany W. Mwaura, James B. Bliska

## Abstract

*Pseudomonas aeruginosa* is an opportunistic bacterial pathogen responsible for a large percentage of airway infections that cause morbidity and mortality in immunocompromised patients, especially those with cystic fibrosis (CF). One important *P. aeruginosa* virulence factor is a type III secretion system (T3SS) that translocates effectors into host cells. ExoS is a T3SS effector with ADP ribosyltransferase (ADPRT) activity. The ADPRT activity of ExoS promotes *P. aeruginosa* virulence by inhibiting phagocytosis and limiting the oxidative burst in neutrophils. The *P. aeruginosa* T3SS also translocates flagellin, which can activate the NLRC4 inflammasome, resulting in: 1) gasdermin-D (GSDMD) pores, release of IL-1β and pyroptosis; and 2) histone 3 citrullination (CitH3) and decondensation and expansion of nuclear DNA into the cytosol. However, recent studies with the *P. aeruginosa* laboratory strain PAO1 indicate that ExoS ADPRT activity inhibits activation of the NLRC4 inflammasome in neutrophils. Here, an ExoS^+^ CF clinical isolate of *P. aeruginosa* with a hyperactive T3SS was identified. Variants of the hyperactive T3SS mutant or PAO1 were used to infect neutrophils from C57BL/6 mice or mice engineered to have a CF genotype or a defect in inflammasome assembly. Responses to NLRC4 inflammasome assembly or ExoS ADPRT activity were assayed, results of which were found to be similar for C57BL/6 or CF neutrophils. The hyperactive T3SS mutant had enhanced resistance to neutrophil killing, like previously identified hypervirulent *P. aeruginosa* isolates. ExoS ADPRT activity in the hyperactive T3SS mutant regulated inflammasome and nuclear DNA decondensation responses like PAO1 but promoted enhanced CitH3 and plasma membrane rupture (PMR). Glycine supplementation inhibited PMR caused by the hyperactive T3SS mutant, suggesting ninjurin-1 is required for this process. These results identify enhanced neutrophil PMR as a pathogenic activity of ExoS ADPRT in a hypervirulent *P. aeruginosa* isolate.

## Introduction

*Pseudomonas aeruginosa* is responsible for a large percentage of airway infections that cause high morbidity and mortality in people with immune deficiencies, such as cystic fibrosis (CF) (1, 2). *P. aeruginosa* has many virulence factors including a type III secretion system (T3SS) that delivers effectors into host cells (3, 4). Three T3SS effectors, ExoT, ExoU and ExoS are important virulence factors (4, 5). All *P. aeruginosa* strains have ExoT, and ExoU or ExoS, but rarely both (6).

ExoS is a bifunctional protein with a Rho GAP domain and an ADP ribosyltransferase (ADPRT) domain (4). The GAP domain targets Rho GTPases, whereas the ADPRT domain modifies multiple proteins, such as Ras (4). The ADPRT domain of ExoS plays a major role in *P. aeruginosa* virulence during infection by blocking phagocytosis and the NADPH oxidase in neutrophils (7–9). ExoS ADPRT activity is important for the PAO1 laboratory strain to survive in neutrophils (8, 10). ExoT is similar to ExoS in structure and also inhibits the NADPH oxidase in neutrophils (8, 10) but is less important for *P. aeruginosa* virulence in airway infections (7, 9).

Infection of neutrophils with T3SS^+^ bacterial pathogens can result in activation of caspase-1 inflammasomes and secretion of the cytokine IL-1β (11). Activation of the NLRC4/caspase-1 inflammasome in neutrophils in response to PAO1 infection and T3SS-mediated secretion of flagellin results in IL-1β secretion (12, 13). Recent studies with PAO1 show that NLRC4/caspase-1 inflammasome activation in neutrophils infected with *P. aeruginosa* can result in IL-1β secretion and cell death by pyroptosis, which requires pore formation by gasdermin D (GSDMD) (14–16). In addition, and somewhat paradoxically, ExoS dampens NLRC4 inflammasome activation (15, 16) and drives IL-1β secretion via NLRP3 (16) in neutrophils infected with PAO1. Neutrophils infected with *exoS^+^* strains like PAO1 can also exhibit signs of NETosis, including citrullination of histone 3 (CitH3), decondensation of nuclear DNA, breakdown of the nuclear membrane, expansion of nuclear DNA into the cytosol, and occasionally extrusion of neutrophil extracellular traps (NETs) (15–17). Santoni et al. reported that most neutrophils infected with PAO1 fail to extrude NETs (incomplete NET extrusion), possibly due to maintenance of the cortical actin cytoskeleton, a process referred to as “incomplete NETosis” (15).

CF is an inherited disease caused by mutations in the CFTR gene which alter movement of chloride and other ions across cell membranes (18). This produces highly viscous mucus in lungs which creates a favorable environment for opportunistic bacterial infection. CF patient mortality is often due to respiratory failure from chronic bacterial infection. ExoS^+^ *P. aeruginosa* strains cause most CF infections (6, 19). The basis for the prevalence of *exoS*^+^ *P. aeruginosa* strains in CF airway infections is unknown and remains an important knowledge gap. *P. aeruginosa* undergoes genetic diversification to generate clonally related populations of variant bacteria during chronic CF airway infections (20). The T3SS may be important for initial colonization of the CF airway, though it is typically inactivated in *P. aeruginosa* during chronic infections (1, 21, 22). However, T3SS function is maintained or hyperactive in some *P. aeruginosa* CF clinical isolates, and this phenotype can be associated with hypervirulence and lung function decline (23, 24). Hyperactive T3SS *P. aeruginosa* strains identified to date contain codon changes *exsD*, which is a negative regulator of the T3SS (23, 24). Two *exoS^+^* CF isolates with hyperactive T3SS phenotypes have codon change mutations T188P or S164P in *exsD* (23, 24). Amplicon sequencing of CF patient samples showed that the frequency of *P. aeruginosa* with the S164P hyperactive *exsD* allele increased during lung function decline, eventually resulting in terminal respiratory failure (23). ExsD binds to and sequesters the transcription factor ExsA (25). The T188P and S164P codon changes in *exsD* are thought to relieve this negative regulation mechanism and result in over secretion of ExoS and ExoT, increased *P. aeruginosa* survival in neutrophils and hypervirulence (23, 24). The outcome of interaction of hyperactive T3SS mutants of *P. aeruginosa* with neutrophils in terms of ExoS-promoted inflammasome modulation, pyroptosis or NETosis has not been reported.

In this study we investigated the outcome of interaction of *P. aeruginosa* with neutrophils in terms of ExoS-promoted inflammasome modulation, pyroptosis and NETosis using PAO1 and CF clinical strains including hyperactive T3SS mutants of *P. aeruginosa.* We studied neutrophils from C57BL/6 mice or isogenic animals engineered to have the most common CFTR mutation (F508del) in the human population. Our results confirm that ExoS activity in PAO1 and clinical strains suppresses the NLRC4 inflammasome (15, 16). In addition, our data indicate that ExoS ADPRT activity promotes plasma membrane rupture (PMR), and this activity, along with bacterial survival, is enhanced with the hyperactive T3SS mutant in C57BL/6 and F508del neutrophils. PMR is inhibited by glycine supplementation, suggesting that ninjurin 1 (NINJ1) (26) is required for this process. In addition, ExoS-promoted PMR was independent of the ASC adaptor, suggesting an inflammasome-independent process. Finally, we find that although ExoS ADPRT promotes generation of CitH3 and PMR, most neutrophils exhibit incomplete NET extrusion, even after infection with the hyperactive T3SS mutant of *P. aeruginosa.* The importance of these findings for understanding the role of ExoS in the pathogenesis of hypervirulent *P. aeruginosa* during neutrophil interactions is discussed.

## Results

### Characterization of BMNs primed with LPS by flow cytometry

For ex vivo infections purified bone marrow neutrophils (BMNs) from C57BL/6 (B6) mice were primed with LPS for ∼18 hr to upregulate inflammasome components (14). Notably, LPS priming can extend neutrophil life spans ex vivo (27, 28). To determine the viability and purity of BMNs, flow cytometry was performed on freshly purified cells as well as purified cells incubated for ∼18 hr with or without 100 ng/ml LPS. BMNs were stained with e780 to assess viability and with neutrophil markers Ly6G and CD11b to measure purity (Supplemental Fig.S1). Dot plots show gating on live cells (e780^-^), and Ly6G^+^ and CD11b^+^ signals (Fig.1). The primed cells (panel C) had a similar percentage of Ly6G^+^ CD11b^+^ as the freshly purified BMNs (panel A). In contrast, BMNs incubated without LPS showed a lower percentage of Ly6G^+^ CD11b^+^ (panel B). These data indicate that our procedure results in relatively pure populations of primed BMNs for infection experiments.

**Fig. 1:**
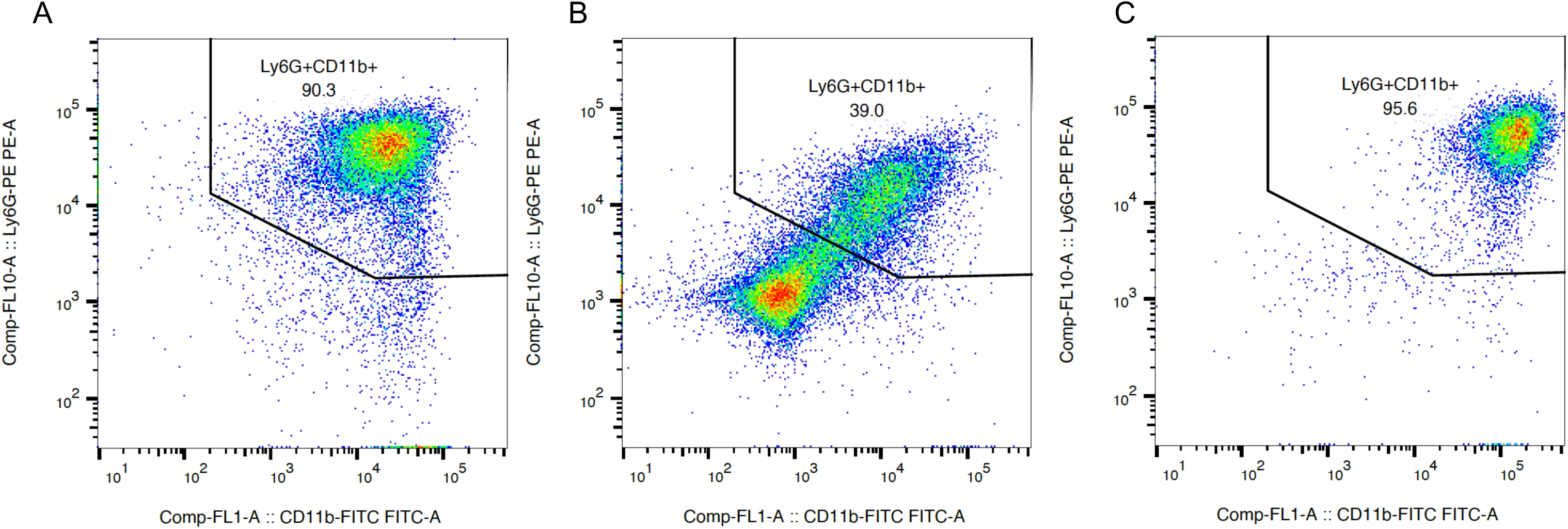
Characterization of BMNs primed with LPS by flow cytometry. BMNs were isolated from B6 mice and analyzed immediately (A) or after ∼18 hr incubation without (B) or with (C) 100 ng/ml LPS. BMNs were stained with e780, Ly6G-PE and CD11b-FITC and analyzed by flow cytometry. Dead cells (e780+) were excluded from the analysis. Representative dot plots of Ly6G and CD11b signals are shown. The gate indicates live cells that are Ly6G^+^ and CD11b^+^.

### ExoS ADPRT activity inhibits IL-1β release in BMNs infected with PAO1

Primed BMNs from B6 mice or those with a Cftr F508del genotype were infected with PAO1 strains (10) to confirm that ExoS ADPRT activity is required for *P. aeruginosa* to inhibit NLRC4 inflammasome activation (16). Results show that more IL-1β was released, as measured by ELISA, from BMNs when they were infected with an ExoS ADPRT catalytic mutant (ExoS(A-)) than the wild type control PAO1F laboratory strain (Fig.2A). As reported by Minns et al. (16) BMNs infected with an ExoT ADPRT catalytic mutant did not release more IL-1β as compared to the control (Supplemental Fig.S2). PMR, as measured by LDH release, also trended higher with ExoS(A-) but was not significant compared to PAO1F (Fig.2B). Additionally, results from CFU assays indicate that the ability of PAO1F to inhibit IL-1β release compared to ExoS(A-) was not due to increased survival of the wild type bacteria during the 60 min infection (Fig.2C). ExoT functions redundantly with ExoS to inhibit the neutrophil NADPH oxidase (8), explaining why ExoS(A-) was not more sensitive to killing than PAO1F. When comparing BMNs from B6 vs. F508del mice there were no major differences between the two in responses to infection with PAO1F or the ExoS(A-) mutant. Together, these data confirm that the ADPRT activity of ExoS but not ExoT is required for PAO1 to inhibit NLRC4 inflammasome activation and IL-1β release in BMNs (16). In addition, the Cftr genotype of the BMNs did not lead to substantial changes in IL-1β release or bactericidal activities.

**Fig. 2.**
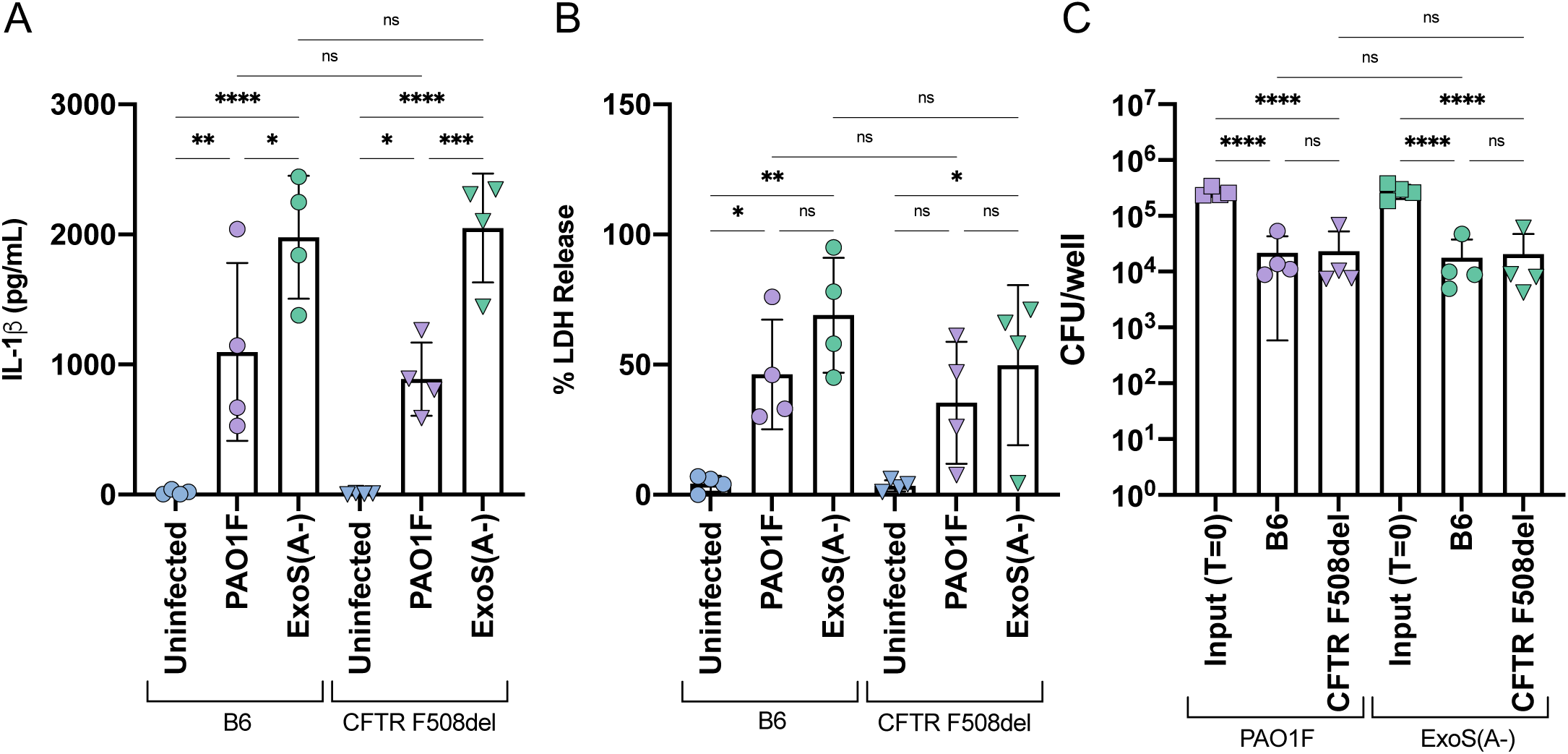
Analysis of BMN infections with PAO1F or ExoS(A-). B6 or CFTR F508del BMNs were left uninfected or infected for 60 min with PAO1F or ExoS(A-) at MOI 10 and analyzed for released IL-1β (A) or LDH (B). (C) BMNs were infected as above at an MOI of 1. Samples of bacteria inoculated into each well (input, T=0) and total bacterial-neutrophil co-cultures solubilized in 0.1% NP-40 detergent (output, T=60m) were processed by serial dilution and plating to determine CFUs. Data represent normalized values for 2.5×10^5^ cells/well (A, B) ± the standard deviation from four independent experiments (A, B, C). Significant differences were determined by two-way ANOVA. ns, not significant; * P<0.05; ** P<0.01; *** P<0.001; **** P<0.0001.

### Identification and characterization of hyperactive T3SS mutant *P. aeruginosa* CF clinical isolates

To increase the biological significance of these results, we studied four clonal isolates that were identified as *exoS^+^ P. aeruginosa* strains that remained T3SS^+^ over time in the sinuses of a CF patient (29). Sinus infections are common in CF patients (29, 30), and they may act as reservoirs that seed *P. aeruginosa* into the lower respiratory tract. Interestingly, two of these isolates, 85 and 86, contained a T48I codon change in *exsA* (29). We analyzed these four isolates for the ability to secrete ExoS *in vitro* (10). Using PAO1F and a T3SS null Δ*pscD* mutant as controls, we found that all strains except Δ*pscD* secreted ExoS, and ExoS was over secreted by 85 and 86 (Fig.3A). These results confirm that patient 32 isolates are T3SS^+^ (29) and suggest that the *exsA*^T48I^ codon change in 85 and 86 is a gain of function mutation leading to the hyperactive T3SS phenotype. To determine if the codon change in *exsA* was sufficient for the ExoS hypersecretion phenotype, allelic exchange was carried out on isolates 8 and 85 to mutate codon 48. A T48I codon change was introduced into 8, and the mutation in 85 was corrected to the wild type sequence. Fig.3B demonstrates that the codon changes in *exsA* reversed the ExoS secretion phenotype in each clinical strain, indicating that the T48I codon change is sufficient to cause the hyperactive T3SS in isolate 85.

**Fig. 3:**
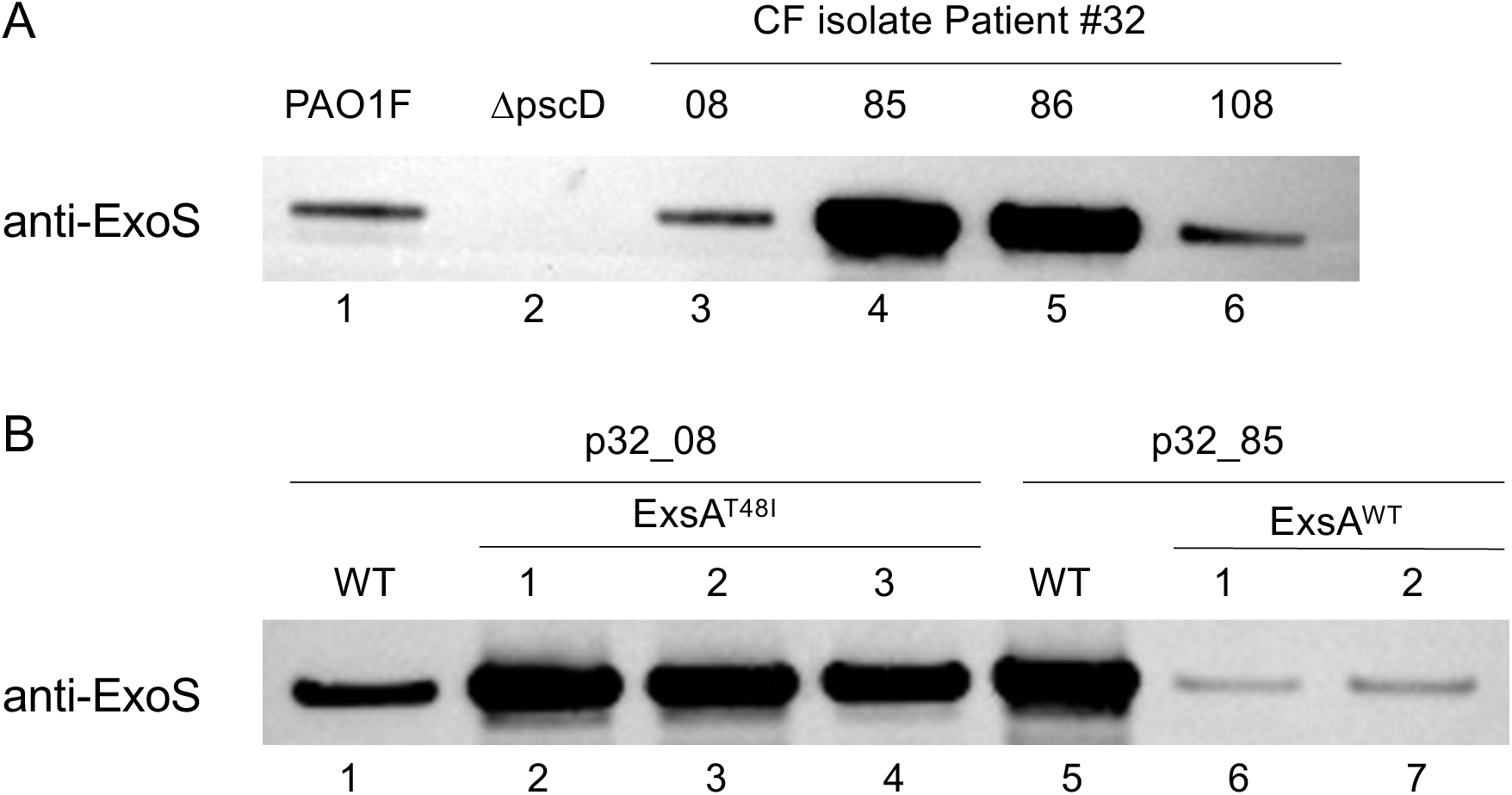
Analysis of ExoS secreted by *P. aeruginosa* isolates. The indicated strains of PAO1F or patient 32 (p32) isolates were grown in bacterial media that activates the T3SS. Secreted ExoS was collected from supernatants and detected by immunoblotting using anti-ExoS antibody. In (B) three independent p32_08 ExsA^T48I^ and two p32_85 ExsA^WT^ mutants were analyzed along with one wild type (WT) control for each.

T48I is in the N-terminal regulatory domain of ExsA that interacts with ExsD (25, 31–33), suggesting that this mutation relieves negative regulation of the T3SS by reducing binding of ExsA to ExsD. To examine this possibility, we used AlphFold2 to predict the structure of the ExsA-ExsD complex, and we highlighted the locations of the T48I mutation, as well as the hyperactivating S164P and T188P codon changes in ExsD (23, 24) (Fig.4). T48 is a solvent exposed residue on a strand of the beta barrel regulatory domain (32) and as shown in Fig.4, is located at the predicted interface of the two proteins, raising the possibility that the T48I change (polar to hydrophobic) interferes with the interaction.

**Fig. 4:**
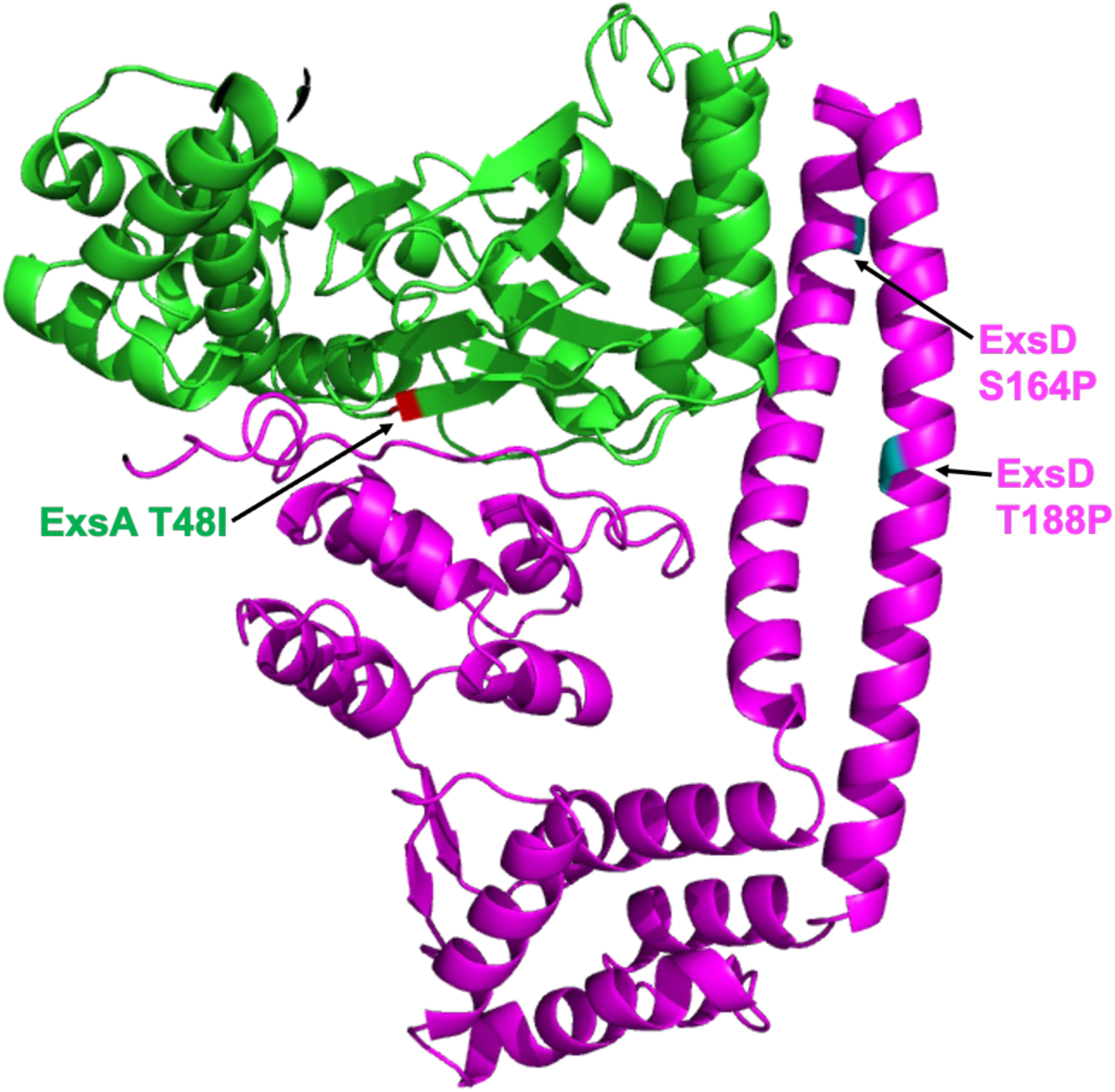
AlphaFold2 prediction of ExsA-ExsD complex and locations of T3SS hyperactivating codon changes. ExsA is shown in green with position of T48I. ExsD is shown in magenta with positions of T188P (Jorth et al. 2015) and S164P (Jorth et al. 2021).

### Hyperactive T3SS mutant *P. aeruginosa* promotes enhanced PMR and reduced bacterial killing in BMNs

We next compared the outcome of infection of BMNs with CF isolates 8 and 85. As shown in Fig.5, levels of IL-1β released were low and similar when BMNs were infected with 8 and 85 (panel A), but 85 stimulated higher LDH release, indicating enhanced PMR, compared to 8 (panel B). In addition, although the clinical isolates 8 and 85 appeared to survive better in BMNs overall as compared to PAO1F (compare with Fig.2), 85 showed significantly higher survival than 8 (Fig.5C). As was seen in Fig. 2 there was no substantial difference when comparing between the B6 and F508del BMNs in their responses to infection (Fig.5).

**Fig. 5:**
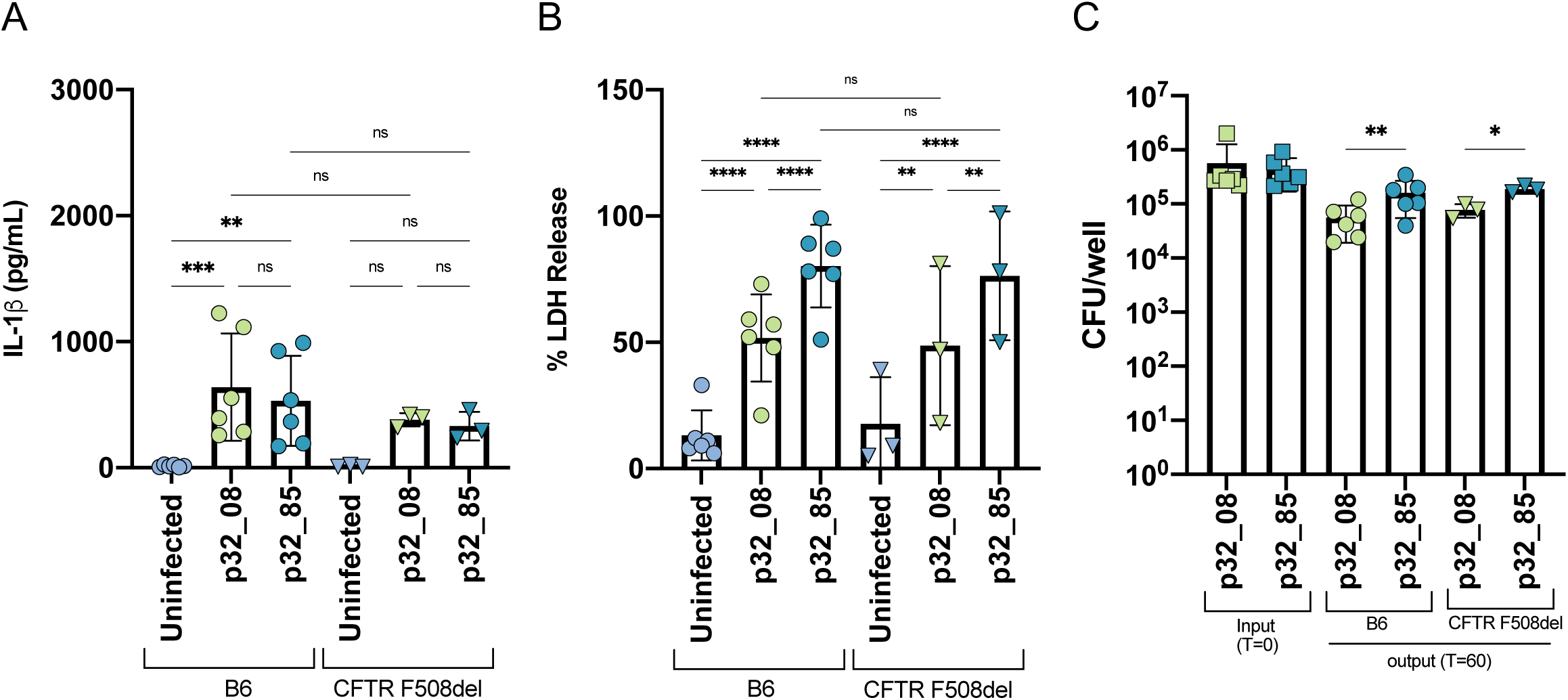
Analysis of BMN infections with patient 32 isolates. B6 or CFTR F508del BMNs were left uninfected or infected for 60 min with p32_8 or p32_85 at MOI 10 (A, B) or 1 (C) and analyzed for released IL-1β (A) or LDH (B) or CFU (C) as in Fig.1. Data represent normalized values for 2.5×10^5^ cells/well (A, B) ± the standard deviation from 3 (CFTR F508del) or 6 (B6) independent experiments (A, B, C). Significant differences were determined by two-way ANOVA. ns, not significant; * P<0.05; ** P<0.01; *** P<0.001; **** P<0.0001.

### Hyperactive T3SS mutant *P. aeruginosa* promotes enhanced CitH3 formation in BMNs

To investigate if ExoS activity and the hyperactive T3SS phenotype impacts NETosis processes we analyzed levels of CitH3 in *P. aeruginosa-*infected BMNs. B6 and F508del BMNs were infected with PAO1F, ExoS(A-), 8 or 85 and immunoblotting was used to measure generation of CitH3 by the calcium regulated enzyme PAD4 (14). Cleavage of GSDMD by caspase-1 was measured as a control since ExoS ADPRT activity inhibits this process (16). As shown in Fig.6A, GSDMD cleavage was highest in B6 BMNs infected with ExoS(A-) as expected (16), although some processing was also detected with PAO1F, 8, and 85, indicating the ExoS block is not complete. CitH3 was generated in response to BMN infection with PAO1F and ExoS(A-), and the elevated production with ExoS(A-) was expected, since NLRC4 activation is associated with this activity (14, 15) (Fig.6A). Interestingly, CitH3 was also elevated with 85 compared to 8 (Fig.6A), suggesting ExoS ADPRT activity in the T3SS hyperactive mutant is driving this process independent of NLRC4 activation. Similar results were obtained with F508del BMNs (Fig.6B), indicating that ExoS ADPRT activity is promoting enhanced CitH3 formation in both CF and non-CF neutrophils infected with the hyperactive T3SS mutant.

**Fig. 6:**
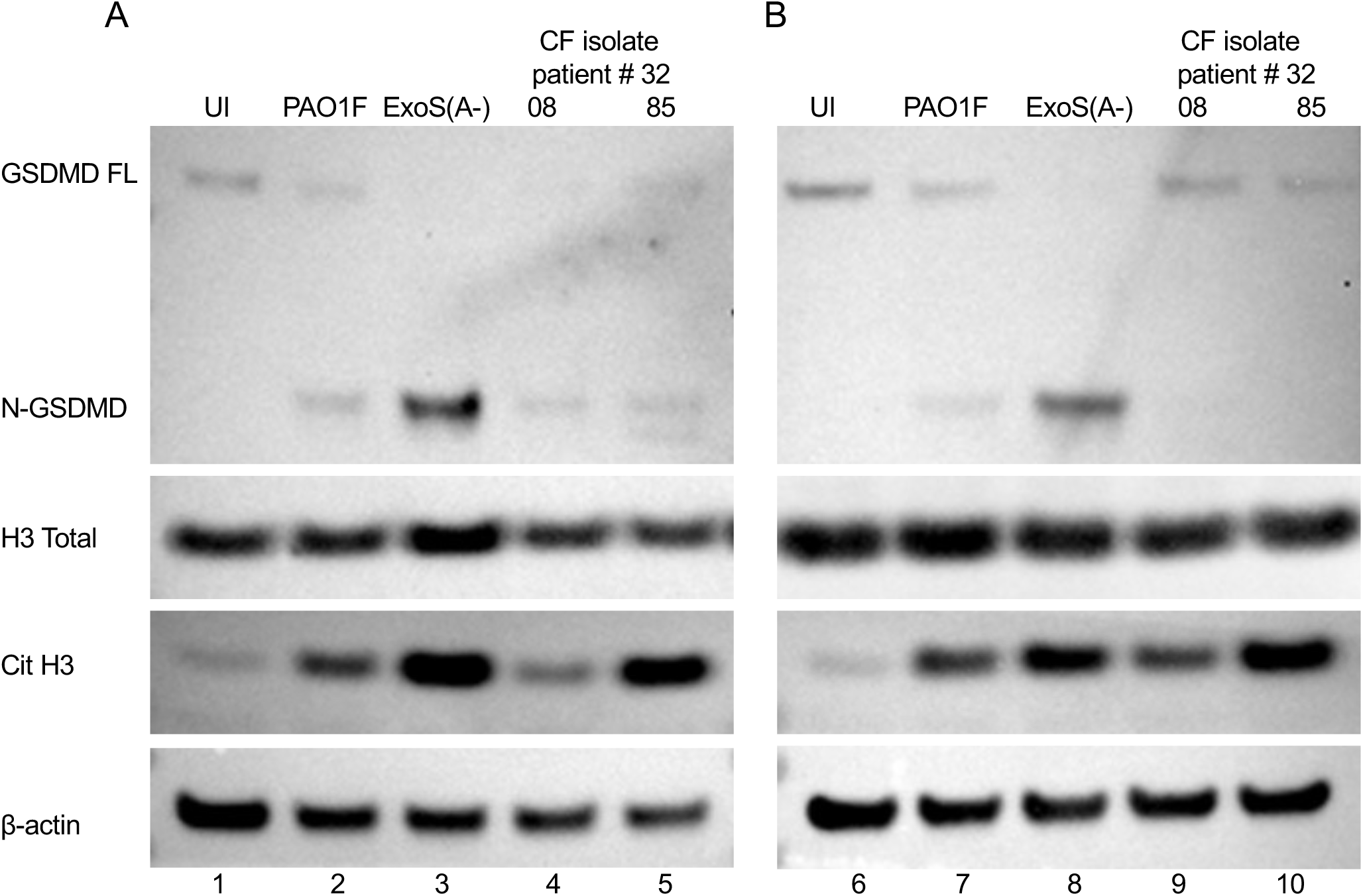
Analysis of BMN infections with PAO1 and patient 32 strains by immunoblotting. BMNs from B6 (A) or F508del (B) mice were left uninfected (UI) or infected with the indicated strains of PAO1F or patient 32 isolates for 60min at an MOI of 10. Samples of total well contents were analyzed by immunoblotting for full length (FL) or cleaved (N-) GSDMD, total histone 3 (H3) and citrullinated histone 3 (Cit H3), and β-actin as a loading control. One representative blot of three independent experiments is shown.

### Hyperactive T3SS mutant *P. aeruginosa* promotes nuclear DNA decondensation and incomplete NET extrusion in BMNs

Since CitH3 is a marker for chromatin decondensation, and a prerequisite for NET extrusion, we wanted to visually determine if infection with PAO1F or our clinical isolates was promoting chromatin decondensation and NET extrusion in BMNs. We first compared PAO1F and PA14, as the latter ExoU^+^ strain was expected to serve as a positive control in microscopy to visualize chromatin decondensation and NET extrusion in B6 BMNs (15). Under our experimental conditions of live cell fluorescence microscopy with the membrane-impermeant DNA dye Sytox Green, we observed with PA14 at 60 min in comparison to uninfected, that there was nuclear DNA decondensation in the majority of BMNs, and a smaller number of cells exhibited extruded NETs (Fig.7A). NET extrusion with PA14 increased at 120 and 180 min as indicated by Sytox Green staining (Fig.7BC). In contrast, in BMNs infected with PAO1F uptake of Sytox Green and DNA decondensation was delayed and even by 180 min these cells produced very few extruded NETs, as compared to PA14 (Fig.7). We next imaged BMNs infected with our additional strains, PAO1F lacking active ExoS ADPRT activity, and both of our clinical isolates. Although we saw increased CitH3 in BMNs infected with ExoS (A-) and the hyperactive T3SS mutant 85 (Fig.6), most cells exhibited only DNA decondensation even at 180 min, and we did not see the robust NET extrusion as with PA14 (Fig.8). These results indicate that hyperactive T3SS mutant *P. aeruginosa,* drives incomplete NET extrusion in BMNs, like the NLRC4-mediated outcome in neutrophils that was observed with an *exoU* mutant (15). Additionally, we observed puncta in the infected cells that could indicate concentrated Sytox Green staining of relaxed chromatin that remained confined within the BMN cell outline (Fig.8).

**Fig. 7:**
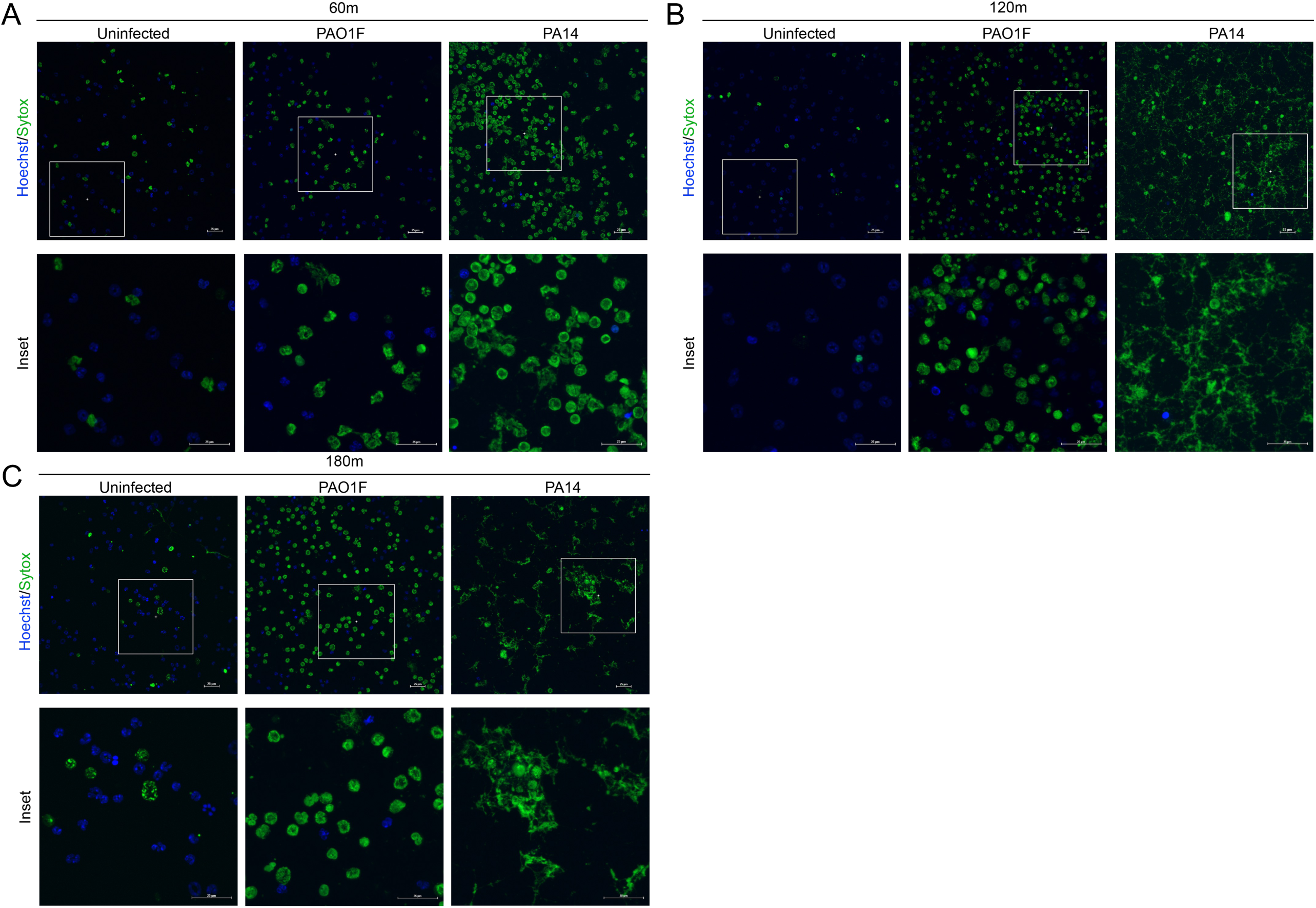
Live cell imaging of B6 neutrophils infected with PAO1F or PA14 Laboratory strains. B6 BMNs were left uninfected or infected for 60 (A), 120 (B), or 180 (C) min with the indicated strains at MOI 10. Cells were stained with Hoechst and Sytox Green 30min prior to imaging with a spinning disc confocal microscope. Images are representative of entire well taken at 40x.

**Fig. 8:**
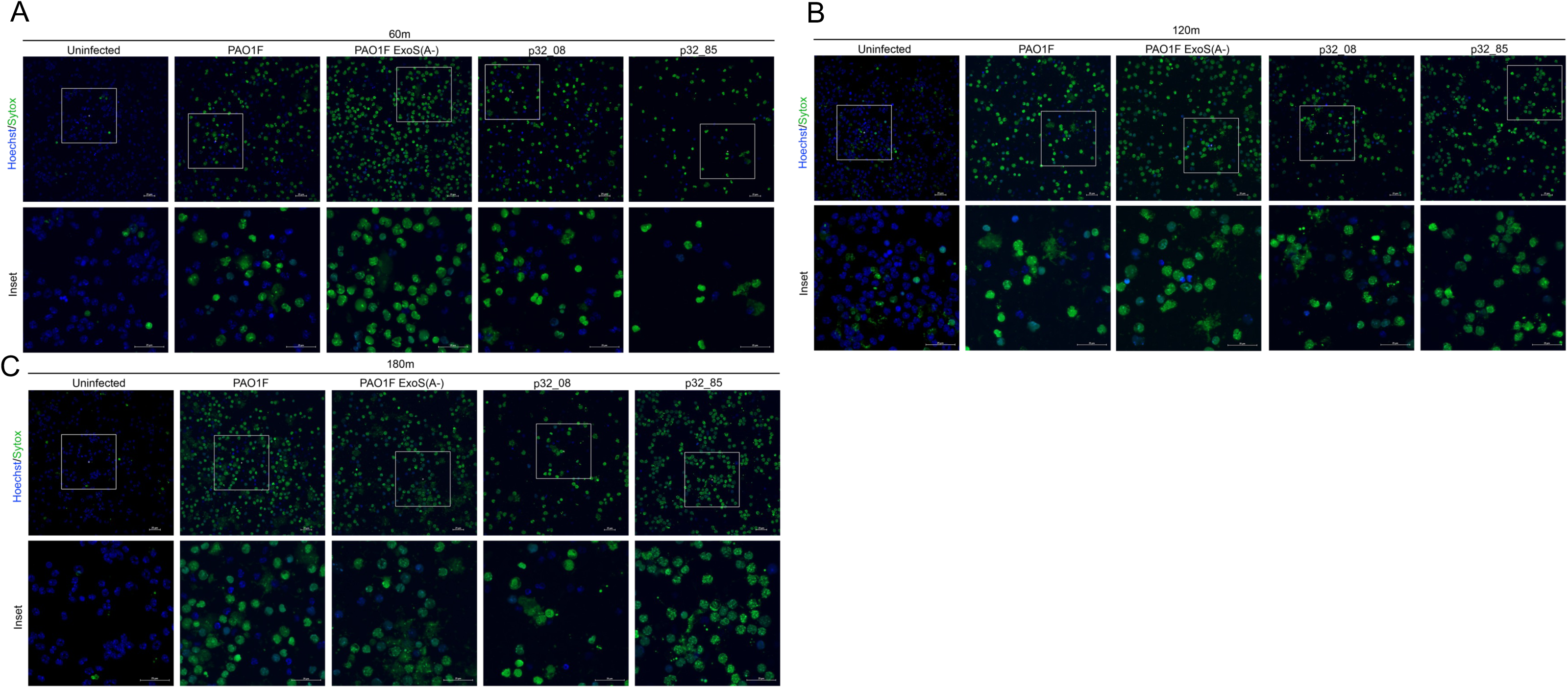
Live Cell Imaging of B6 Neutrophils infected with PAO1F or patient 32 isolates. B6 BMNs were left uninfected or infected for 60 (A), 120 (B), or 180 (C) min with indicated strains at MOI 10. Cells were stained with Hoechst and Sytox Green 30min prior to imaging by spinning disc confocal microscopy. Images are representative of entire wells taken at 40x.

### ASC contributes to IL-1β release but is dispensable for enhanced PMR in BMNs infected with hyperactive T3SS mutant *P. aeruginosa*

When evaluating potential mechanisms for the enhanced PMR in BMNs infected with hyperactive T3SS mutant *P. aeruginosa* we considered the possible role of inflammasome activation. Although ExoS suppresses the NLRC4 inflammasome (15, 16), cleavage of GSDMD and release of IL-1β is not completely blocked (Figs.2,6), and Minns et al. obtained evidence that ExoS ADPRT activity drives activation of NLRP3 in BMNs infected with PAO1 (16). To examine the role of inflammasome activation in PMR we used Asc^-/-^ BMNs, which lack the adaptor that is required for, or can contribute to, NLRP3 and NLRC4 inflammasome assembly, respectively (34). B6 or Asc^-/-^ BMNs were infected with PAO1F, ExoS(A-), 8 or 85 and LDH release was determined to measure PMR. We also measured secreted IL-1β and immunoblotted for GSDMD cleavage and CitH3. Fig.9B shows that PMR was not significantly reduced for any of the strains comparing B6 to Asc^-/-^ BMNs. Strain 85 continued to cause enhanced PMR in Asc^-/-^ BMNs as compared to 8 (Fig.9B). Notably, IL-1β release was significantly reduced for all strains in the absence of ASC (Fig.9A). There were corresponding decreases in cleavage of GSDMD in the absence of ASC for all strains (Fig.9C). These results indicate a role for ASC in the NCRC4 inflammasome triggered by ExoS(A-) infection, and the NLRP3 inflammasome driven by ExoS ADPRT activity (16) with the other strains (Fig.9A). Interestingly, CitH3 production was reduced in the absence of ASC for ExoS(A-) but not 85 (Fig.9C). Together, these finding suggests that ExoS ADPRT-promoted PMR and CitH3 production in response to infection with the hyperactive T3SS mutant is inflammasome-independent.

**Fig. 9:**
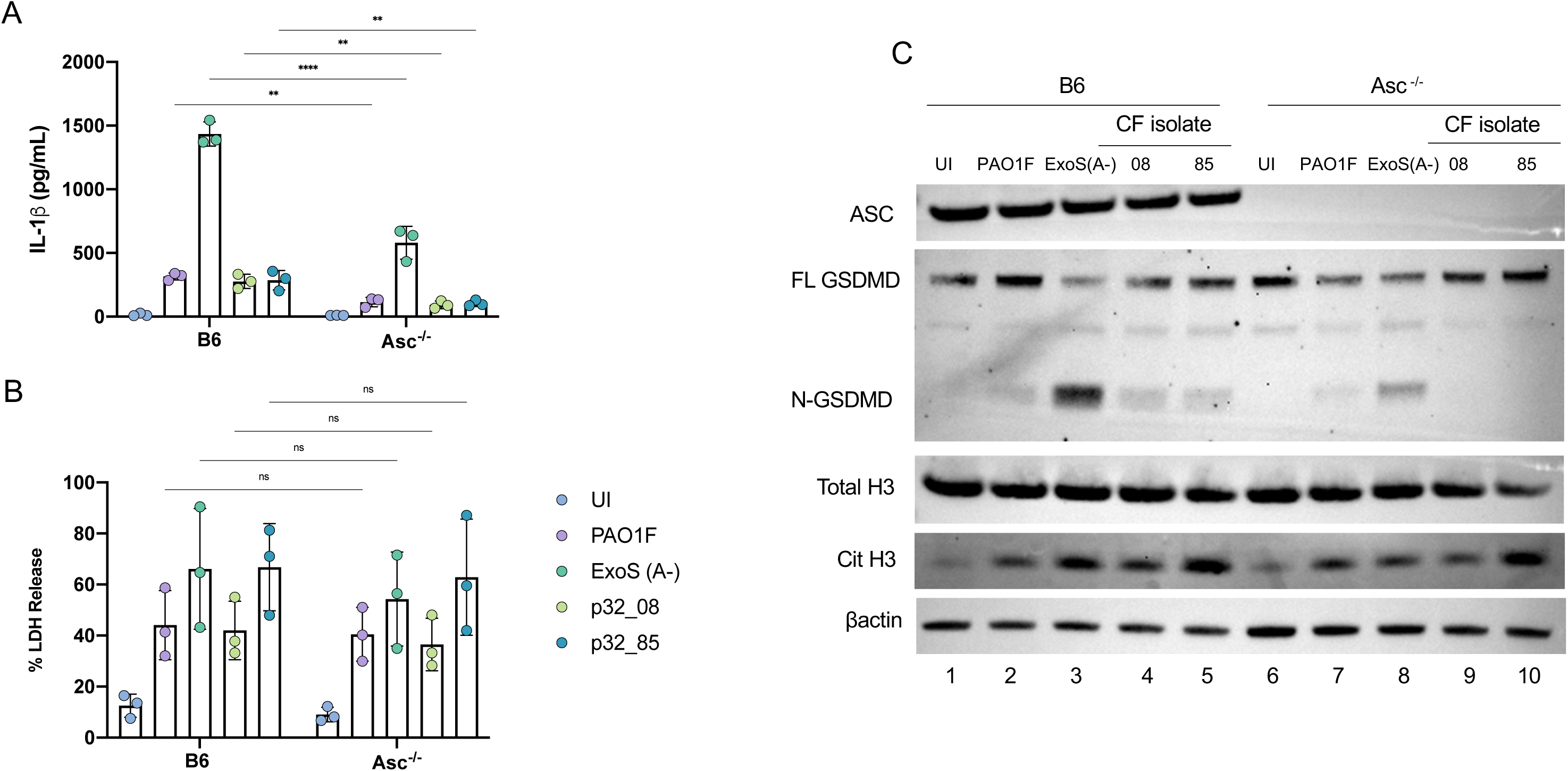
Analysis of B6 and ASC^-/-^ BMN infections with PAO1 or patient 32 strains. B6 or *ASC^-/-^* BMNs were left uninfected or infected for 60 min with indicated strains at MOI 10 and analyzed for released IL-1β (A) or LDH (B). Data represent normalized values for 2.5×10^5^ cells/well ± the standard deviation (IL-1β, LDH) from three independent experiments. (C) Samples of total well contents were analyzed by immunoblotting for ASC, total histone 3 (H3) or citrullinated histone 3 (Cit H3), full length (FL) or cleaved (N-) GSDMD, and β-actin as a loading control. One representative blot of three independent experiments is shown. (A,B) Significant differences were determined by two-way ANOVA compared to ExoS(A-) or between groups. ns, not significant; * P<0.05; **; **** P<0.0001.

### Glycine inhibits PMR promoted by ExoS ADPRT activity in BMNs infected with hyperactive T3SS mutant *P. aeruginosa*

Recently a new mechanism for PMR has been identified, in which the NINJ1 protein oligomerizes downstream of different cell death pathways, leading to the formation of large lesions in the plasma membrane (35, 36). PMR has been shown to be inhibited by the addition of the amino acid glycine, which is thought to prevent the oligomerization of NINJ1 (26). Santoni et al. found that glycine selectively inhibited PMR but not IL-1β release in BMNs infection with a PAO1 Δ*exoS* mutant, demonstrating a selective cytoprotective effect against NINJ1 lesion formation but not GSDMD pore formation downstream of NLRC4 inflammasome activation (15). We tested a role for NINJ1 in PMR in our BMN model by adding 5mM glycine to the media during *P. aeruginosa* infection. BMNs were infected with the clinical strains 8 or 85, as well as their corresponding ADPRT catalytic activity mutants ExoS(A-) constructed by allelic exchange. LDH release was determined to measure PMR, and as controls we also measured secreted IL-1β and immunoblotted for GSDMD cleavage and CitH3. Indeed, we found that glycine significantly reduced LDH release in all infections, indicating a reduction in NINJ1-mediated PMR (Fig.10B). In contrast, glycine addition had no impact on cleavage of GSDMD (Fig.10C) or release of IL-1β (Fig.10A) in the infected BMNs. The use of the ExoS(A-) mutants confirmed that ADPRT activity is required for ExoS in the patient 32 strains to inhibit the NLRC4 inflammasome (Fig.10A,C). Additionally, we saw no difference in the level of CitH3 between glycine conditions (Fig.10C). Together, these results suggest that NINJ1 can mediate PMR in BMNs in response to either NLRC4 inflammasome activation (15) or ExoS ADPRT activity (Fig. 10). In addition, either NLRC4 inflammasome activation or ExoS ADPRT activity can promote formation of CitH3 independent of NINJ1-mediated PMR (Fig.10). In the case of the former, GSDMD pores may promote CitH3 (14) and NINJ1 lesions (15). The type of membrane perturbation that promotes CitH3 generation and activation of NINJ1 in response to ExoS ADPRT activity is unknown.

**Fig. 10:**
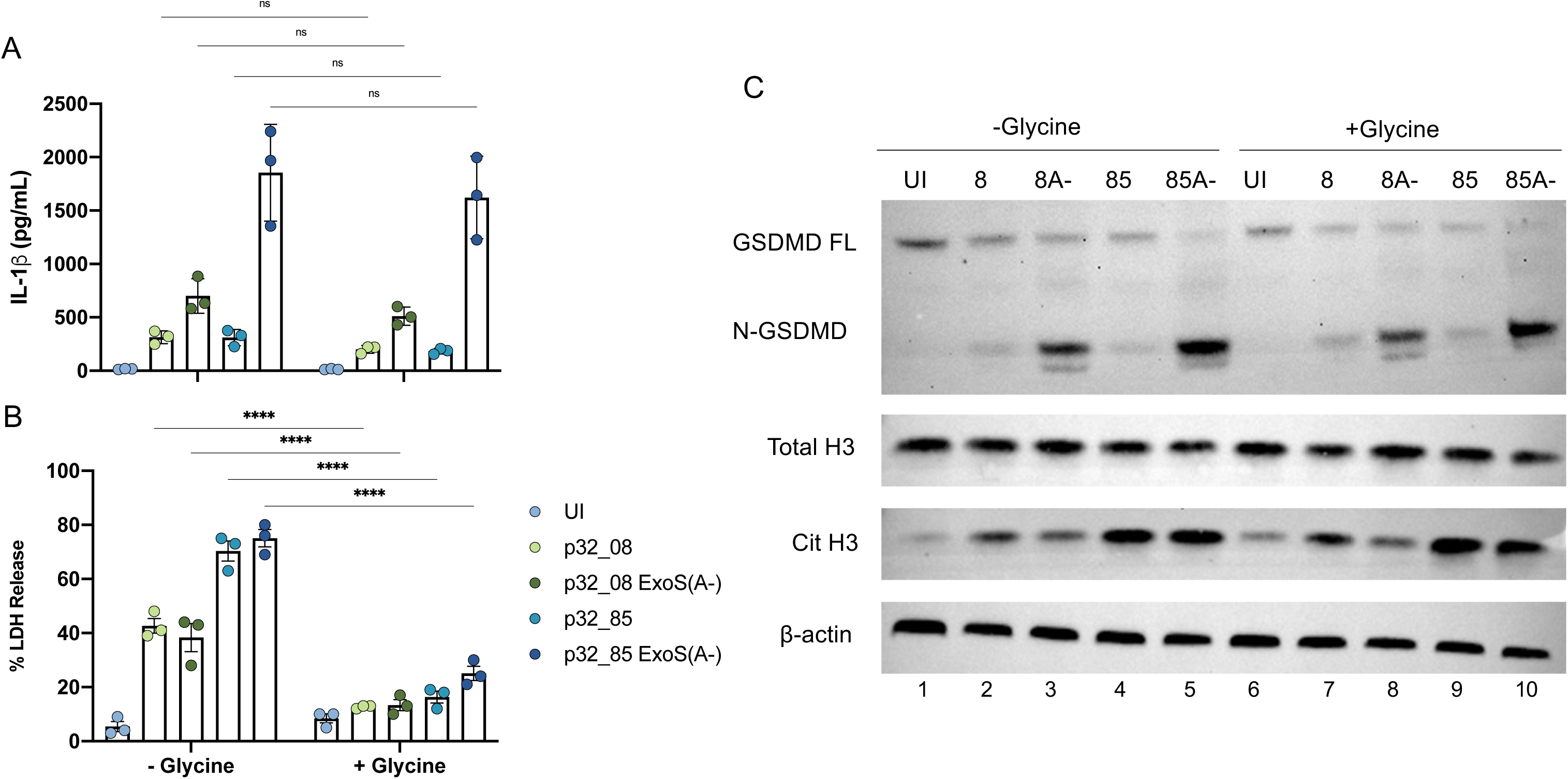
Analysis of B6 BMNs infections with patient 32 strains in absence or presence of glycine. B6 BMNs were left uninfected or infected for 60 min with indicated strains at MOI 10 in the absence or presence of 5 mM glycine and analyzed for released IL-1β (A) or LDH (B). Data represent normalized values for 2.5×10^5^ cells/well ± the standard deviation (IL-1β, LDH) from three independent experiments. (C) Samples of total well contents were analyzed by immunoblotting for total histone 3 (H3), citrullinated histone 3 (Cit H3), Gasdermin D (GSDMD), and β-actin as a loading control. One representative blot of three independent experiments is shown. (A, B) Significant differences were determined by two-way ANOVA between treatment groups. ns, not significant; **** P<0.0001.

## Discussion

Results presented here are consistent with recent reports indicating that upon infection of neutrophils with PAO1 the *P. aeruginosa* T3SS translocates flagellin, which activates the NLRC4 inflammasome, resulting in GSDMD pores, release of IL-1β, pyroptosis, generation of CitH3 and decondensation of chromatin into incomplete NETs, and activation of NINJ1 leading to PMR (12, 14–16). In addition, our data are in line with the idea that the ADPRT activity of translocated ExoS dampens activation of NLRC4 and promotes IL-1β release via the NLRP3 inflammasome in neutrophils infected with PAO1 (15, 16). We identified an ExoS^+^ *P. aeruginosa* hyperactive T3SS mutant clinical strain from a CF infection with enhanced resistance to neutrophil bactericidal activity, like the hypervirulent isolates identified by Jorth et al. (23, 24). The *P. aeruginosa* hyperactive T3SS mutant was used to obtain new evidence that ExoS ADPRT activity promotes CitH3, and activation of NINJ1 leading to PMR in CF and non-CF murine neutrophils.

The clonal hyperactive T3SS mutants with a T48I codon change in *exsA* were identified by genome sequencing of individual colonies obtained at a single time point (day 553) in a longitudinal study of a CF patient with a chronic *P. aeruginosa* sinus infection (29). Sinuses are thought to act as reservoirs for generating *P. aeruginosa* mutants with enhanced fitness that seed the lower respiratory tract (29). Unfortunately, no data is available to suggest increased airway disease in patient 32 at day 553 or if the hyperactive T3SS mutant successfully colonized the lungs (29). It is also unclear if *exsA^T48I^* was the majority allele in the *P. aeruginosa* population in the sinuses at day 553, or if it was coincidental that the colonies selected contained this mutation. It is possible that patient 32 was fortunate in avoiding dissemination of *exsA^T48I^* from the sinuses, since the two known *exsD* hyperactive T3SS mutants that have been isolated were associated with serious lung function decline in CF patients (23, 24). Going forward it will be important to determine if and how ExsA T48I reduces interaction of ExsA with ExsD. The ExsD S164P and T188P codon variants are also likely defective for binding to ExsA (23, 24), as the Pro insertions could disrupt the alpha helices that are in proximity to ExsA (Fig. 4). Defining how these codon changes impact ExsA and ExsD functions will provide additional insight into the interaction mechanism of these two proteins. It will also be interesting to determine if hypervirulent *P. aeruginosa* strains are unknowingly present in strain collections from diverse infections sources, as this would increase the clinical significance of these isolates.

Our experiments prioritize ExoS over ExoT because: first, the former is more important for *P*. *aeruginosa* virulence in airway infections (7, 9); second, the *exoS*^+^ gene is epidemiologically linked to successful CF airway infection by *P*. *aeruginosa* (6, 19); and third ExoS but not ExoT is required to inhibit GSDMD cleavage and IL-1β release from BMNs (15, 16). In addition, unlike ExoS, ExoT is not required for LDH release in BMNs lacking caspase-1 (Casp1^-/-^) (15), indicating that ExoT does not contribute to PMR. However, ExoT ADPRT activity can independently reduce NADPH oxidase-dependent bactericidal activity against PAO1 in neutrophils (8, 10) and the same is likely true for hypervirulent *P. aeruginosa* strains.

The ExoS ADPRT domain has broad specificity for host protein targets making it challenging to determine the mechanism by which it dampens activation of NLRC4 and promotes IL-1β release via the NLRP3 inflammasome, generation of CitH3 and decondensation of chromatin into incomplete NETS, and activation of NINJ1 leading to PMR in murine neutrophils. In addition, it is striking that ExoS does not suppress NLRC4 or activate the NLRP3 inflammasome in murine bone marrow-derived macrophages infected with PAO1 (16). Minns et al. suggest that ExoS ADPRT inhibits NAIPs/NLRC4 and activates NLRP3 in murine neutrophils by direct modifications, and that increases in the levels of these substrates in murine macrophages makes these cells less sensitive to intoxication (16). Minns et al. obtained evidence that the priming state of murine neutrophils may also lead to different responses to PAO1 infections due to impacts on levels of inflammasome components and ExoS substrates (16). It is also possible that ExoS ADPRT activity indirectly inhibits NAIPs/NLRC4 and activates NLRP3, while simultaneously promoting generation of CitH3 and decondensation of chromatin into incomplete NETS, and activation of NINJ1 leading to PMR in neutrophils. An early study reported that ExoS ADPRT activity promotes apoptosis in HeLa cells infected with *P. aeruginosa* (37). Experiments using inhibitors of capases-8 and −3 could be used to investigate the role of apoptosis pathways in neutrophil responses to ExoS ADPRT activity.

Santoni et al. presented preliminary evidence that ExoS contributes to PMR in murine or human neutrophils infected with PAO1 (15), but additional experiments are needed to define if ADPRT activity dampens activation of NLRC4, generation of CitH3 and decondensation of chromatin into incomplete NETS, and activation of NINJ1 leading to PMR in human cells. It will also be important to determine if there is any impact of CF genotype on neutrophil response to ExoS ADPRT activity. Thus far we have not observed major phenotypic differences between non-CF and CF BMNs. However, differences between non-CF and CF human neutrophils have been reported in terms of antimicrobial responses and NET extrusion (38) and therefore these comparisons are important.

Pad4 is a calcium-dependent enzyme and evidence suggests that Ca^2+^ influx through GSDMD pores is needed for CitH3 generation in response to NLRC4 inflammasome activation in infected neutrophils (14, 15). Our results suggest that enhanced CitH3 generation in response to infection with hyperactive T3SS mutant *P. aeruginosa* can occur in the absence of detectable GSDMD cleavage in Asc^-/-^ neutrophils. ExoS ADPRT activity in neutrophils infected with hyperactive T3SS mutant *P. aeruginosa* may be promoting Ca^2+^ influx by some other mechanism, leading enhanced CitH3 generation. Alternatively, it is possible that GSDMD pore formation below our limit of detection allows Ca^2+^ influx under these conditions. The use of Casp1^-/-^ BMNs to further decrease the possibility of GSDMD pore formation would help address these possibilities.

Our results using glycine supplementation with neutrophils suggest that PMR promoted by ExoS ADPRT activity is dependent on activation and oligomerization of NINJ1, resulting in large lesions in the plasma membrane (26, 35, 36). How NINJ1 is activated in response to membrane perturbations is unknown. Formation of GSDMD pores appears to be one mechanism leading to NINJ1 activation. Degen et al. suggest that NINJ1 senses changes in membrane composition, such as the exposure of negatively charged phosphatidylserine on the plasma membrane (35). We considered the possibility that NINJ1-mediated plasma membrane damage was involved in Ca^2+^ influx needed for enhanced CitH3 generation in response to infection with hyperactive T3SS mutant *P. aeruginosa*. However, enhanced CitH3 generation was not reduced by glycine supplementation. This result may indicate that glycine only prevents NINJ1 oligomerization to form large lesions but does not prevent Ca^2+^ influx. Alternatively, ExoS ADPRT activity may promote membrane perturbations that in turn allow Ca^2+^ influx and NINJ1 activation.

Although the mechanism by which ExoS ADPRT activity promotes generation of CitH3 and decondensation of chromatin into incomplete NETS, and activation of NINJ1 leading to PMR in neutrophils is unknown, these responses could represent cell intrinsic mechanisms of bactericidal or pathogenic activities impacting *P. aeruginosa.* Santoni et al. compared survival of PAO1 in BMNs that were deficient in Pad4 (Padi4^-/-^) or NLRC4 (Nlrc4^-/-^) or control wild type (15). Results indicate that generation of CitH3 and incomplete NET extrusion under control of Pad4 had no impact on bacterial survival, while Nlrc4^-/-^ BMNs were significantly more bactericidal (15). These results suggest that a neutrophil response under control of NLRC4, such as PMR, is pathogenic. Future experiments will be needed to determine if ExoS-promoted PMR favors survival of *P. aeruginosa* during neutrophil infection. It would be informative to measure survival of hyperactive T3SS mutant *P. aeruginosa* in Ninj1^-/-^ neutrophils to determine if enhanced PMR contributes to the hypervirulent phenotype of these strains.

## Acknowledgements

This work was funded by the Dartmouth Cystic Fibrosis Training Program (T32 HL134598), P30 DK117469 from the NIH and the Cystic Fibrosis Foundation (STANTO19R0).

We acknowledge the important contributions of Natasha Medici to initiating study of ExoS and ExoT in the laboratory. We would like to thank Arne Rietsch and Jennifer Bomberger for kindly providing us with *P. aeruginosa* strains, and Joshua Obar for Asc^-/-^ mice. We would also like to thank Ko-Wei Liu and Alexander Rapp for assistance with acquisition and analysis of flow cytometry data.

We would also like to thank the Dartmouth Life Sciences Light Microscopy facility which is supported by bioMT through NIH grant P20-GM113132.

We acknowledge Dartmouth Genomics and Molecular Biology Shared Resources Core Facility (NCI Cancer Center, Support Grant 5P30CA023108-37). Flow cytometry experiments were carried out in DartLab, the Immune Monitoring and Flow Cytometry Shared Resource at the Norris Cotton Cancer Center at Dartmouth, with NCI Cancer Center Support Grant 5P30 CA023108-41 and COBRE grant P30GM103415-15.

## Author Contributions

A.D.R. designed/performed experiments, analyzed data, generated figures and wrote the manuscript. B.W.M designed and created alphafold2 model of ExsA-ExsD co-crystal structure. J.B.B. conceived/funded the study, designed experiments, and wrote the manuscript.

## Materials and Methods

### Bacterial Strains

Bacterial strains used in this study are outlined in Table 1. *P. aeruginosa* strains for neutrophil infections and secretion assays were grown on Luria-Bertani (LB) agar plates, and in LB high salt broth (11.7g/L NaCl) supplemented with 0.2mM MgCl_2_ and 0.5M CaCl_2_ at 37°C (10). *P. aeruginosa* for strain construction were grown in LB broth. *Escherichia coli* for strain construction were grown in LB supplemented with the appropriate antibiotic.

**Table 1:** Strains and Plasmids.

| Strain or Plasmid | Relevant genotype or description | Reference or Source |
| --- | --- | --- |
| **Strains** |  |  |
| 1560 (RP 1951(RP 1831)) | PAO1F, wild type | Arne Rietsch |
| 1561 (RP 1903) | PAO1F $\Delta pscD$ | Arne Rietsch |
| 1697 (RP 5481) | PAO1F *exoS*(A-) | Arne Rietsch |
| 1698 (RP 6202) | PAO1F *exoT*(A-) | Arne Rietsch |
| 1699 (RP 6205) | PAO1F *exoS*(A-) *exoT*(A-) | Arne Rietsch |
| 1725 | CF isolate patient 32 #8 | Jennifer Bomberger |
| 1726 | CF isolate patient 32 #85 | Jennifer Bomberger |
| 1727 | CF isolate patient 32 #86 | Jennifer Bomberger |
| 1728 | CF isolate patient 32 #108 | Jennifer Bomberger |
| 1771 | CF isolate patient 32 #8 *exoS*(A-) | This study |
| 1781 | CF isolate patient 32 #85 *exoS*(A-) | This study |
| 1757 | CF isolate patient 32 #8 *exsA*<sup>T48I</sup> | This study |
| 1760 | CF isolate patient 32 #85 *exsA*<sup>I48T(WT)</sup> | This study |
| **Plasmids** |  |  |
| pTOPO2.1 | Cloning vector | Invitrogen |
| pMQ30 | Allelic exchange vector | George O'Toole |

### Plasmids

Primer sequences used to amplify and mutagenize ExoS and ExsA are outlined in Table 2. The ExoS and ExsA genes were amplified from patient 32 isolates 8 and 85 using standard colony PCR and cloned into pTOPO2.1 and transformed into *E. coli* DH5α. Site directed mutagenesis was performed on pTOPO-ExoS to introduce two point mutations, E379D and E381D, to inactivate ExoS ADPRT catalytic activity following the protocol provided by Agilent Technologies. All constructs were then cloned from pTOPO into pMQ30 using standard restriction digestion and ligation and transformed into DH5α for long term storage and S17 *E. coli* for conjugation with *P. aeruginosa*.

**Table 2:**
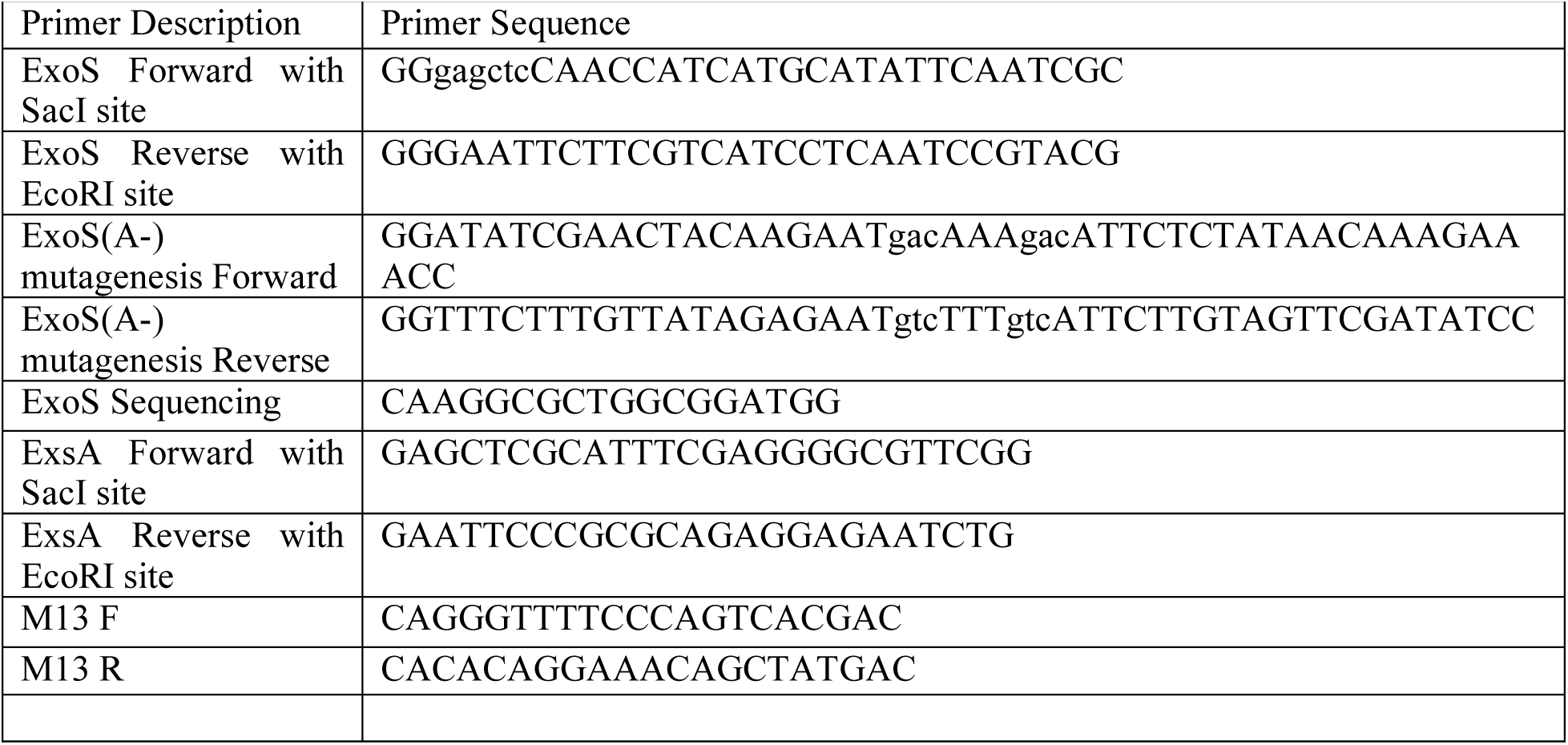
Primers.

### Conjugation and Allelic Exchange

Strains of *P. aeruginosa* for conjugation were grown overnight at 42°C without shaking in LB media. *E. coli* strains carrying the allelic exchange vector were grown overnight at 30°C with shaking in LB with appropriate antibiotic. Constructs were introduced into *P. aeruginosa* by conjugation with S17-1 *E. coli.* Merodiploids were selected by antibiotic resistance. Double recombinants were acquired using 10% sucrose counter selection (39). PCR and sanger sequencing was used to confirm mutations.

### Ethics Statement

Isolation of bone marrow from mice was carried out in accordance with a protocol that adheres to the Guide for the Care and Use of Laboratory Animals of the National Institutes of Health (NIH) and was reviewed and approved (approval 2148) by the Institutional Animal Care and Use Committee at Dartmouth College. The Dartmouth College animal program is registered with the U.S. Department of Agriculture (USDA) through certificate number 12-R-0001, operates in accordance with Animal Welfare Assurance (NIH/PHS) under assurance number D16-00166 (A3259-01) and is accredited with the Association for Assessment and Accreditation of Laboratory Animal Care International (AAALAC, accreditation number 398).

### Mouse Strains

C57BL06J (stock#664) mice were purchased from Jackson Laboratories. Mice with the *Cftr^F508del^* mutation on the C57BL/6 background were obtained from Case Western Reserve University’s Cystic Fibrosis Mouse Models Core and bred at Dartmouth (40). CCR2-GFP^+^ (41) were bred and housed at Dartmouth college. Asc^-/-^ mice (42) were acquired from the lab of Joshua Obar at Dartmouth College. CCR2-GFP^+^ BMNs were used as wild type controls for Asc^-/-^ BMNs in the experiments shown in Fig.9. All mice used in this study were in the C57BL/6 genetic background and were bred in and housed up to four mice per ventilated cage at the Dartmouth CCMR facility unless otherwise stated. Both male and female mice aged between 8-12 weeks were used for neutrophil isolation experiments.

### Preparation of Neutrophils and Flow Cytometry

Bone marrow was isolated from tibia and femur exudates of 8-12 week old C57BL/6J (Jackson Laboratories), CCR2-eGFP^+^, ASC^-/-^ or *Cftr^F508del^*. Single-cell suspensions of bone marrow were prepared, and neutrophils were isolated following Miltenyi MACS Neutrophil Isolation Kit. Neutrophils were plated at a density of 2.5×10^5^ in 100μL of OptiMEM in a 96 well plate. The neutrophils were primed with 100ng/mL O26:B6 *Escherichia coli* LPS (Sigma) and incubated overnight (16h) at 37°C with 5% CO_2_. Purity was assessed by flow-cytometry, staining for live cells using e780, and CD11b and Ly6G for neutrophils. Cell fluorescence was detected by Beckman Coulter Cytoflex S and analyzed using FlowJo software.

### Neutrophil Infection Assays

Overnight (16-hour) cultures of *P. aeruginosa* were sub-cultured 1:100 in fresh LB high-salt broth supplemented with 10mM MgCl_2_ and 0.5mM CaCl_2_ and 5mM EGTA and grown at 37°C to mid-log phase at OD_600_= 0.5. Cultures were then pelleted, the LB removed, and bacteria were resuspended in PBS to the original volume. Bacterial suspensions were then diluted to an MOI of 10 and 1 in OptiMEM. Overnight media from the neutrophils was removed and replaced with 100μL of OptiMEM containing bacteria. Plates were then incubated at 37°C with 5% CO_2_ for 60, 120, or 180 minutes. Cell supernatants (MOI 10) were collected for cytokine ELISAs and lactate dehydrogenase (LDH) assays. Whole wells (MOI 10) were lysed with 1x sample buffer (Invitrogen) containing DTT (Invitrogen), complete mini (Roche) protease inhibitor and PhosSTOP (Roche) phosphatase inhibitor. Cell supernatants were removed (MOI 1), and cells were lysed with 100μL 0.1% NP-40 and serially diluted in PBS and plated on LB agar for CFU assays.

### ExoS secretion assay

Overnight (16-hour) cultures of *P. aeruginosa* were sub-cultured 1:100 in fresh LB high-salt broth supplemented with 10mM MgCl_2_ and 0.5mM CaCl_2_ and 5mM EGTA and grown at 37°C to mid-log phase at OD_600_= 0.5. 1ml of culture was pelleted, supernatant was mixed with Trichloro Acetic Acid (final concentration 10%) and incubated overnight at 4°C with shaking. Proteins were pelleted by centrifugation, washed with acetone and resuspended in 1x sample buffer containing DTT. Secreted proteins were resolved on SDS page gel and western blotted for ExoS.

### Western blotting

Cell lysates were run on 4-12% NuPAGE Bis-Tris SDS-PAGE gels (Invitrogen by ThermoFisher Scientific) and transferred to PVDF membranes (ThermoFisher Scientific) using an iBlot 2 Gel Transfer Device (Life Technologies). Membranes were blocked in 5% non-fat dairy milk and incubated with primary antibody overnight. The primary antibodies used were rabbit MAb for GSDMD (Abcam, ab209845), MAb for Histone 3 (Cell Signaling #96C10), Mab for Citrullinated Histone 3 (Abcam #ab5103) and rabbit polyclonal for b-actin (Cell Signaling, #4967), and polyclonal ab for ExoS (from Arne Rietsch). HRP-conjugated anti-rabbit (Jackson Immuno Research) or HRP-conjugated anti-mouse (Jackson Immuno Research) was used as a secondary antibody. Protein bands reacting with antibodies were visualized using chemiluminescent detection reagent (GE Healthcare) on an iBright FL1500 (ThermoFisher Scientific).

### IL-1β quantification

Murine IL-1b in neutrophils supernatants was quantified using ELISA kits (R&D Systems^®^, MLB00C) following the manufacturer’s instructions.

### LDH quantification

LDH in neutrophil supernatants was quantified using the CytoTox 96® Non-Radioactive Cytotoxicity Assay (Promega®) following the manufacturer’s instructions.

### Visualization of nuclear DNA decondensation and incomplete NET Extrusion

BMNs were seeded into glass bottom 96-well plates at a density of 2.5 x 10^5^ cells/well. Overnight (16h) primed BMNs were infected with *P. aeruginosa* at an MOI of 10 for 60, 120, or 180 minutes. DNA was detected by cell impermeable DNA fluorescent dye (2.5μΜ Sytox Green) along with cell-permeable DNA dye (1μg/ml Hoechst). Dyes were added 30 minutes before imaging. Fluorescent images were taken with a 40x objective with a Nikon spinning disc confocal microscope. Images were processed and analyzed using the Nikon NIS-Elements AR software (version 5.42.02).

### ExsA-ExsD complex modeling

A model of the tertiary structure of the ExSA-ExSD complex was generated using the Alphafold2-multimer tool of ColabFold (43). The protein sequences of ExSA and ExSD were entered into the Alphafold2-multimer server and run using default settings. PyMOL was used to generate illustrative figures and label the hyperactivating codon changes.

### Statistical analysis of data

Experimental data generated from at least three independent in vitro experiments are analyzed for significance using GraphPad Prism software. Probability (*p*) values for ELISA and LDH data are calculated using one-way analysis of variance (ANOVA) or grouped two-way ANOVA with Tukey’s posttest. P values of ≤ 0.05 were considered significant (*p ≤ 0.05, **p ≤ 0.01, ***p ≤ 0.001, ns: not significant).

---

**Fig. S1:**
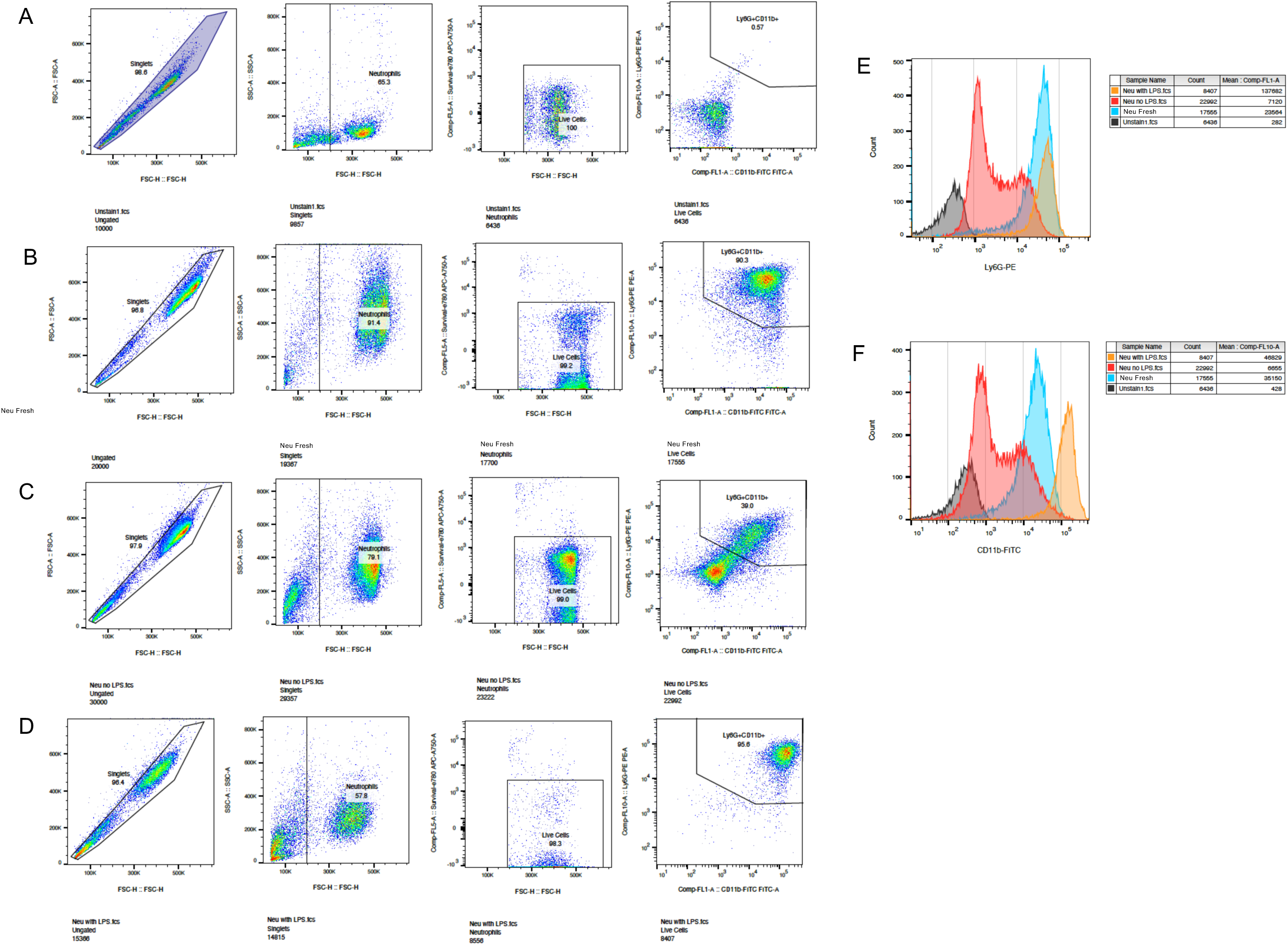
Characterization of BMNs primed with LPS by flow cytometry. BMNs were isolated from B6 mice and analyzed immediately without staining (A), immediately with D staining (B) or after 18 hr incubation without (C) or with (D) 100 ng/ml LPS. BMNs were stained with e780, Ly6G-PE and CD11b-FITC and analyzed by flow cytometry. (A-D) The gates indicate from left to right: singlets, normal size, live cells, and cells that are Ly6G^+^ and CD11b^+^. Histograms show overlay of fluorescence intensity for Ly6G^+^ (E) or CD11b^+^ (F) in each group.

**Fig. S2.**
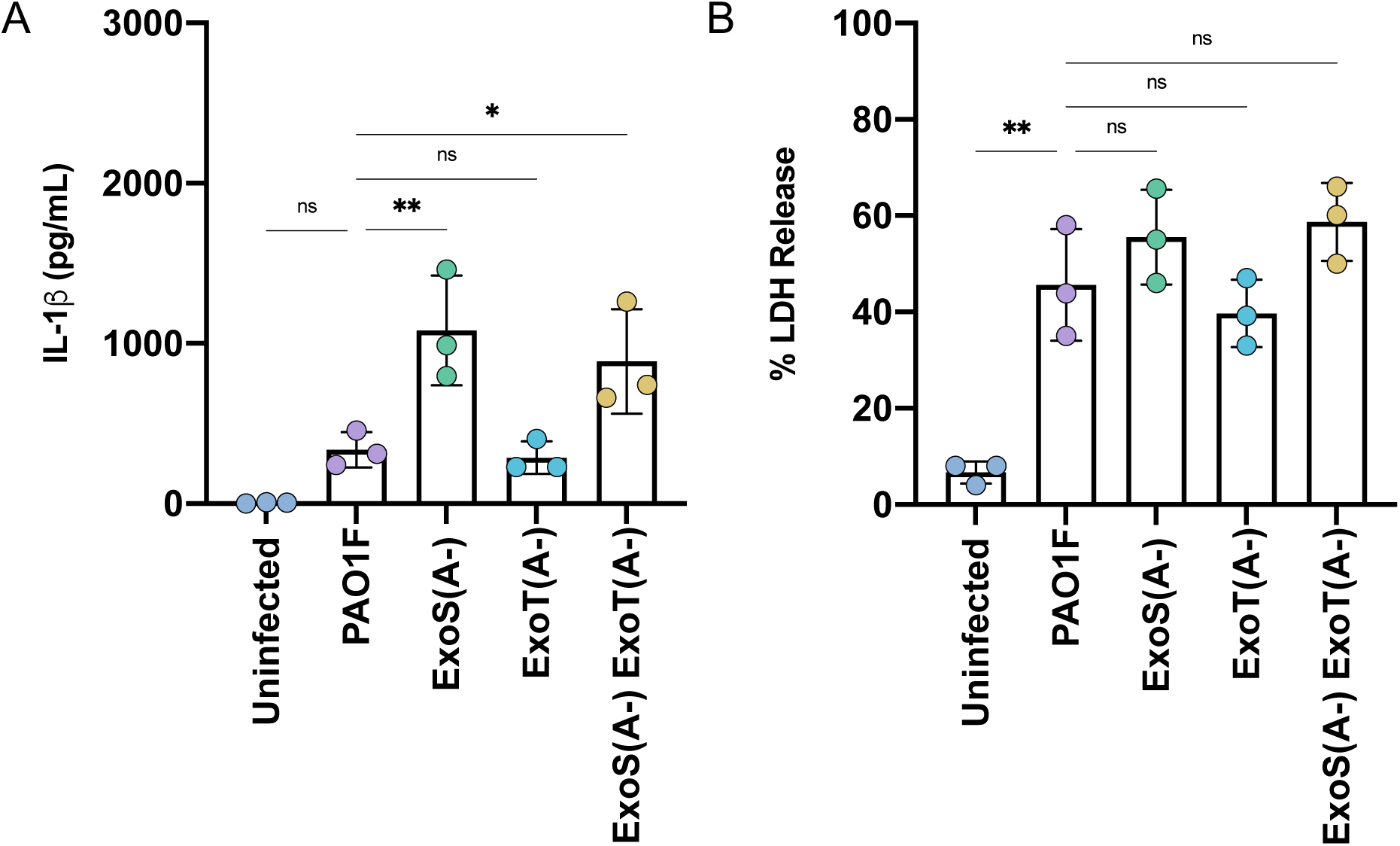
Analysis of BMN infections with PAO1F or ADPRT catalytic mutants. B6 BMNs were left uninfected or infected for 60 min with PAO1F, ExoS(A-), ExoT(A-), or the double ExoS(A-)ExoT(A-) double mutant at MOI 10 and analyzed for released IL-1β (A) or LDH (B). Data represent normalized values for 2.5×10^5^ cells/well ± the standard deviation from four independent experiments. Significant differences were determined by two-way ANOVA. ns, not significant; * P<0.05; ** P<0.01;.

## References

1. Hauser AR, Jain M, Bar-Meir M, McColley SA. Clinical significance of microbial infection and adaptation in cystic fibrosis. Clin Microbiol Rev. 2011;24(1):29–70. Epub 2011/01/15. doi: 10.1128/CMR.00036-10. PubMed PMID: 21233507; PMCID: PMC3021203.

2. Malhotra S, Hayes D, Jr., Wozniak DJ. Cystic Fibrosis and Pseudomonas aeruginosa: the Host-Microbe Interface. Clin Microbiol Rev. 2019;32(3). Epub 2019/05/31. doi: 10.1128/CMR.00138-18. PubMed PMID: 31142499; PMCID: PMC6589863.

3. Galle M, Carpentier I, Beyaert R. Structure and function of the Type III secretion system of Pseudomonas aeruginosa. Curr Protein Pept Sci. 2012;13(8):831–42. Epub 2013/01/12. doi: 10.2174/138920312804871210. PubMed PMID: 23305368; PMCID: PMC3706959.

4. Hauser AR. The type III secretion system of Pseudomonas aeruginosa: infection by injection. Nat Rev Microbiol. 2009;7(9):654–65. Epub 2009/08/15. doi: 10.1038/nrmicro2199. PubMed PMID: 19680249; PMCID: PMC2766515.

5. Mohamed MF, Gupta K, Goldufsky JW, Roy R, Callaghan LT, Wetzel DM, Kuzel TM, Reiser J, Shafikhani SH. CrkII/Abl phosphorylation cascade is critical for NLRC4 inflammasome activity and is blocked by Pseudomonas aeruginosa ExoT. Nat Commun. 2022;13(1):1295. Epub 2022/03/13. doi: 10.1038/s41467-022-28967-5. PubMed PMID: 35277504; PMCID: PMC8917168.

6. Feltman H, Schulert G, Khan S, Jain M, Peterson L, Hauser AR. Prevalence of type III secretion genes in clinical and environmental isolates of Pseudomonas aeruginosa. Microbiology (Reading). 2001;147(Pt 10):2659–69. Epub 2001/09/29. doi: 10.1099/00221287-147-10-2659. PubMed PMID: 11577145.

7. Rangel SM, Logan LK, Hauser AR. The ADP-ribosyltransferase domain of the effector protein ExoS inhibits phagocytosis of Pseudomonas aeruginosa during pneumonia. mBio. 2014;5(3):e01080–14. Epub 2014/06/12. doi: 10.1128/mBio.01080-14. PubMed PMID: 24917597; PMCID: PMC4056551.

8. Vareechon C, Zmina SE, Karmakar M, Pearlman E, Rietsch A. Pseudomonas aeruginosa Effector ExoS Inhibits ROS Production in Human Neutrophils. Cell Host Microbe. 2017;21(5):611–8 e5. Epub 2017/05/12. doi: 10.1016/j.chom.2017.04.001. PubMed PMID: 28494242; PMCID: PMC5478421.

9. Shaver CM, Hauser AR. Relative contributions of Pseudomonas aeruginosa ExoU, ExoS, and ExoT to virulence in the lung. Infect Immun. 2004;72(12):6969–77. Epub 2004/11/24. doi: 10.1128/IAI.72.12.6969-6977.2004. PubMed PMID: 15557619; PMCID: PMC529154.

10. Sun Y, Karmakar M, Taylor PR, Rietsch A, Pearlman E. ExoS and ExoT ADP ribosyltransferase activities mediate Pseudomonas aeruginosa keratitis by promoting neutrophil apoptosis and bacterial survival. J Immunol. 2012;188(4):1884–95. Epub 2012/01/18. doi: 10.4049/jimmunol.1102148. PubMed PMID: 22250085; PMCID: PMC3273577.

11. Yow SJ, Yeap HW, Chen KW. Inflammasome and gasdermin signaling in neutrophils. Mol Microbiol. 2022;117(5):961–72. Epub 2022/03/05. doi: 10.1111/mmi.14891. PubMed PMID: 35244299.

12. Ryu JC, Kim MJ, Kwon Y, Oh JH, Yoon SS, Shin SJ, Yoon JH, Ryu JH. Neutrophil pyroptosis mediates pathology of P. aeruginosa lung infection in the absence of the NADPH oxidase NOX2. Mucosal Immunol. 2017;10(3):757–74. Epub 2016/08/25. doi: 10.1038/mi.2016.73. PubMed PMID: 27554297.

13. Huus KE, Joseph J, Zhang L, Wong A, Aaron SD, Mah TF, Sad S. Clinical Isolates of Pseudomonas aeruginosa from Chronically Infected Cystic Fibrosis Patients Fail To Activate the Inflammasome during Both Stable Infection and Pulmonary Exacerbation. J Immunol. 2016;196(7):3097–108. Epub 2016/02/21. doi: 10.4049/jimmunol.1501642. PubMed PMID: 26895832.

14. Oh C, Li L, Verma A, Reuven AD, Miao EA, Bliska JB, Aachoui Y. Neutrophil inflammasomes sense the subcellular delivery route of translocated bacterial effectors and toxins. Cell Rep. 2022;41(8):111688. Epub 2022/11/24. doi: 10.1016/j.celrep.2022.111688. PubMed PMID: 36417874.

15. Santoni K, Pericat D, Gorse L, Buyck J, Pinilla M, Prouvensier L, Bagayoko S, Hessel A, Leon-Icaza SA, Bellard E, Mazeres S, Doz-Deblauwe E, Winter N, Paget C, Girard JP, Pham CTN, Cougoule C, Poincloux R, Lamkanfi M, Lefrancais E, Meunier E, Planes R. Caspase-1-driven neutrophil pyroptosis and its role in host susceptibility to Pseudomonas aeruginosa. PLoS Pathog. 2022;18(7):e1010305. Epub 2022/07/19. doi: 10.1371/journal.ppat.1010305. PubMed PMID: 35849616; PMCID: PMC9345480.

16. Minns MS, Liboro K, Lima TS, Abbondante S, Miller BA, Marshall ME, Tran Chau J, Roistacher A, Rietsch A, Dubyak GR, Pearlman E. NLRP3 selectively drives IL-1beta secretion by Pseudomonas aeruginosa infected neutrophils and regulates corneal disease severity. Nat Commun. 2023;14(1):5832. Epub 2023/09/21. doi: 10.1038/s41467-023-41391-7. PubMed PMID: 37730693; PMCID: PMC10511713.

17. Skopelja-Gardner S, Theprungsirikul J, Lewis KA, Hammond JH, Carlson KM, Hazlett HF, Nymon A, Nguyen D, Berwin BL, Hogan DA, Rigby WFC. Regulation of Pseudomonas aeruginosa-Mediated Neutrophil Extracellular Traps. Front Immunol. 2019;10:1670. Epub 2019/08/06. doi: 10.3389/fimmu.2019.01670. PubMed PMID: 31379861; PMCID: PMC6657737.

18. Ratjen F, Bell SC, Rowe SM, Goss CH, Quittner AL, Bush A. Cystic fibrosis. Nat Rev Dis Primers. 2015;1:15010. Epub 2015/01/01. doi: 10.1038/nrdp.2015.10. PubMed PMID: 27189798.

19. Ozer EA, Nnah E, Didelot X, Whitaker RJ, Hauser AR. The Population Structure of Pseudomonas aeruginosa Is Characterized by Genetic Isolation of exoU+ and exoS+ Lineages. Genome Biol Evol. 2019;11(1):1780–96. Epub 2019/06/08. doi: 10.1093/gbe/evz119. PubMed PMID: 31173069; PMCID: PMC6690169.

20. Williams D, Evans B, Haldenby S, Walshaw MJ, Brockhurst MA, Winstanley C, Paterson S. Divergent, coexisting Pseudomonas aeruginosa lineages in chronic cystic fibrosis lung infections. Am J Respir Crit Care Med. 2015;191(7):775–85. Epub 2015/01/16. doi: 10.1164/rccm.201409-1646OC. PubMed PMID: 25590983; PMCID: PMC4407486.

21. Jain M, Bar-Meir M, McColley S, Cullina J, Potter E, Powers C, Prickett M, Seshadri R, Jovanovic B, Petrocheilou A, King JD, Hauser AR. Evolution of Pseudomonas aeruginosa type III secretion in cystic fibrosis: a paradigm of chronic infection. Transl Res. 2008;152(6):257–64. Epub 2008/12/09. doi: 10.1016/j.trsl.2008.10.003. PubMed PMID: 19059160; PMCID: PMC2628760.

22. Banwart B, Splaingard ML, Farrell PM, Rock MJ, Havens PL, Moss J, Ehrmantraut ME, Frank DW, Barbieri JT. Children with cystic fibrosis produce an immune response against exoenzyme S, a type III cytotoxin of Pseudomonas aeruginosa. J Infect Dis. 2002;185(2):269–70. Epub 2002/01/25. doi: 10.1086/338197. PubMed PMID: 11807706.

23. Jorth P, Durfey S, Rezayat A, Garudathri J, Ratjen A, Staudinger BJ, Radey MC, Genatossio A, McNamara S, Cook DA, Aitken ML, Gibson RL, Yahr TL, Singh PK. Cystic Fibrosis Lung Function Decline after Within-Host Evolution Increases Virulence of Infecting Pseudomonas aeruginosa. Am J Respir Crit Care Med. 2021;203(5):637–40. Epub 2020/11/03. doi: 10.1164/rccm.202007-2735LE. PubMed PMID: 33137262; PMCID: PMC7924579.

24. Jorth P, Staudinger BJ, Wu X, Hisert KB, Hayden H, Garudathri J, Harding CL, Radey MC, Rezayat A, Bautista G, Berrington WR, Goddard AF, Zheng C, Angermeyer A, Brittnacher MJ, Kitzman J, Shendure J, Fligner CL, Mittler J, Aitken ML, Manoil C, Bruce JE, Yahr TL, Singh PK. Regional Isolation Drives Bacterial Diversification within Cystic Fibrosis Lungs. Cell Host Microbe. 2015;18(3):307–19. Epub 2015/08/25. doi: 10.1016/j.chom.2015.07.006. PubMed PMID: 26299432; PMCID: PMC4589543.

25. Brutinel ED, Vakulskas CA, Yahr TL. Functional domains of ExsA, the transcriptional activator of the Pseudomonas aeruginosa type III secretion system. J Bacteriol. 2009;191(12):3811–21. Epub 2009/04/21. doi: 10.1128/JB.00002-09. PubMed PMID: 19376850; PMCID: PMC2698394.

26. Borges JP, Saetra RSR, Volchuk A, Bugge M, Devant P, Sporsheim B, Kilburn BR, Evavold CL, Kagan JC, Goldenberg NM, Flo TH, Steinberg BE. Glycine inhibits NINJ1 membrane clustering to suppress plasma membrane rupture in cell death. Elife. 2022;11. Epub 2022/12/06. doi: 10.7554/eLife.78609. PubMed PMID: 36468682; PMCID: PMC9754625.

27. Parker LC, Whyte MK, Dower SK, Sabroe I. The expression and roles of Toll-like receptors in the biology of the human neutrophil. J Leukoc Biol. 2005;77(6):886–92. Epub 2005/02/25. doi: 10.1189/jlb.1104636. PubMed PMID: 15728244.

28. Miralda I, Uriarte SM, McLeish KR. Multiple Phenotypic Changes Define Neutrophil Priming. Front Cell Infect Microbiol. 2017;7:217. Epub 2017/06/15. doi: 10.3389/fcimb.2017.00217. PubMed PMID: 28611952; PMCID: PMC5447094.

29. Armbruster CR, Marshall CW, Garber AI, Melvin JA, Zemke AC, Moore J, Zamora PF, Li K, Fritz IL, Manko CD, Weaver ML, Gaston JR, Morris A, Methe B, DePas WH, Lee SE, Cooper VS, Bomberger JM. Adaptation and genomic erosion in fragmented Pseudomonas aeruginosa populations in the sinuses of people with cystic fibrosis. Cell Rep. 2021;37(3):109829. Epub 2021/10/24. doi: 10.1016/j.celrep.2021.109829. PubMed PMID: 34686349; PMCID: PMC8667756.

30. Robertson JM, Friedman EM, Rubin BK. Nasal and sinus disease in cystic fibrosis. Paediatr Respir Rev. 2008;9(3):213–9. Epub 2008/08/13. doi: 10.1016/j.prrv.2008.04.003. PubMed PMID: 18694713.

31. Brutinel ED, Vakulskas CA, Yahr TL. ExsD inhibits expression of the Pseudomonas aeruginosa type III secretion system by disrupting ExsA self-association and DNA binding activity. J Bacteriol. 2010;192(6):1479–86. Epub 2009/12/17. doi: 10.1128/JB.01457-09. PubMed PMID: 20008065; PMCID: PMC2832532.

32. Shrestha M, Bernhards RC, Fu Y, Ryan K, Schubot FD. Backbone Interactions Between Transcriptional Activator ExsA and Anti-Activator ExsD Facilitate Regulation of the Type III Secretion System in Pseudomonas aeruginosa. Sci Rep. 2020;10(1):9881. Epub 2020/06/20. doi: 10.1038/s41598-020-66555-z. PubMed PMID: 32555263; PMCID: PMC7303211.

33. Shrestha M, Xiao Y, Robinson H, Schubot FD. Structural Analysis of the Regulatory Domain of ExsA, a Key Transcriptional Regulator of the Type Three Secretion System in Pseudomonas aeruginosa. PLoS One. 2015;10(8):e0136533. Epub 2015/09/01. doi: 10.1371/journal.pone.0136533. PubMed PMID: 26317977; PMCID: PMC4552939.

34. Patankar YR, Mabaera R, Berwin B. Differential ASC requirements reveal a key role for neutrophils and a noncanonical IL-1beta response to Pseudomonas aeruginosa. Am J Physiol Lung Cell Mol Physiol. 2015;309(8):L902-13. doi: 10.1152/ajplung.00228.2015. PubMed PMID: 26472815; PMCID: PMC4609944.

35. Degen M, Santos JC, Pluhackova K, Cebrero G, Ramos S, Jankevicius G, Hartenian E, Guillerm U, Mari SA, Kohl B, Muller DJ, Schanda P, Maier T, Perez C, Sieben C, Broz P, Hiller S. Structural basis of NINJ1-mediated plasma membrane rupture in cell death. Nature. 2023;618(7967):1065-71. Epub 2023/05/18. doi: 10.1038/s41586-023-05991-z. PubMed PMID: 37198476; PMCID: PMC10307626.

36. Kayagaki N, Kornfeld OS, Lee BL, Stowe IB, O’Rourke K, Li Q, Sandoval W, Yan D, Kang J, Xu M, Zhang J, Lee WP, McKenzie BS, Ulas G, Payandeh J, Roose-Girma M, Modrusan Z, Reja R, Sagolla M, Webster JD, Cho V, Andrews TD, Morris LX, Miosge LA, Goodnow CC, Bertram EM, Dixit VM. NINJ1 mediates plasma membrane rupture during lytic cell death. Nature. 2021;591(7848):131-6. Epub 2021/01/21. doi: 10.1038/s41586-021-03218-7. PubMed PMID: 33472215.

37. Kaufman MR, Jia J, Zeng L, Ha U, Chow M, Jin S. Pseudomonas aeruginosa mediated apoptosis requires the ADP-ribosylating activity of exoS. Microbiology (Reading). 2000;146 (Pt 10):2531–41. Epub 2000/10/06. doi: 10.1099/00221287-146-10-2531. PubMed PMID: 11021928.

38. Robledo-Avila FH, Ruiz-Rosado JD, Brockman KL, Kopp BT, Amer AO, McCoy K, Bakaletz LO, Partida-Sanchez S. Dysregulated Calcium Homeostasis in Cystic Fibrosis Neutrophils Leads to Deficient Antimicrobial Responses. J Immunol. 2018;201(7):2016–27. Epub 2018/08/19. doi: 10.4049/jimmunol.1800076. PubMed PMID: 30120123; PMCID: PMC6143431.

39. Mould DL, Stevanovic M, Ashare A, Schultz D, Hogan DA. Metabolic basis for the evolution of a common pathogenic Pseudomonas aeruginosa variant. Elife. 2022;11. Epub 20220503. doi: 10.7554/eLife.76555. PubMed PMID: 35502894; PMCID: PMC9224983.

40. Loeven NA, Perault AI, Cotter PA, Hodges CA, Schwartzman JD, Hampton TH, Bliska JB. The Burkholderia cenocepacia Type VI Secretion System Effector TecA Is a Virulence Factor in Mouse Models of Lung Infection. mBio. 2021;12(5):e0209821. Epub 2021/09/29. doi: 10.1128/mBio.02098-21. PubMed PMID: 34579569; PMCID: PMC8546862.

41. Serbina NV, Hohl TM, Cherny M, Pamer EG. Selective expansion of the monocytic lineage directed by bacterial infection. J Immunol. 2009;183(3):1900–10. Epub 2009/07/15. doi: 10.4049/jimmunol.0900612. PubMed PMID: 19596996; PMCID: PMC2753883.

42. Mariathasan S, Newton K, Monack DM, Vucic D, French DM, Lee WP, Roose-Girma M, Erickson S, Dixit VM. Differential activation of the inflammasome by caspase-1 adaptors ASC and Ipaf. Nature. 2004;430(6996):213-8. Epub 2004/06/11. doi: 10.1038/nature02664. PubMed PMID: 15190255.

43. Mirdita M, Schutze K, Moriwaki Y, Heo L, Ovchinnikov S, Steinegger M. ColabFold: making protein folding accessible to all. Nat Methods. 2022;19(6):679–82. Epub 2022/06/01. doi: 10.1038/s41592-022-01488-1. PubMed PMID: 35637307; PMCID: PMC9184281.

